# Systemic Metabolic Depletion of Intestine Microbiome Undermines Melanoma Immunotherapy Effectiveness

**DOI:** 10.1101/2023.10.09.561540

**Authors:** Natalia V. Zakharevich, Maxim D. Morozov, Vera A. Kanaeva, Artem B. Ivanov, Vladimir I. Ulyantsev, Ksenia M. Klimina, Evgenii I. Olekhnovich

**Affiliations:** Lopukhin Federal Research and Clinical Center of Physical-Chemical Medicine of Federal Medical Biological Agency, Malaya Pirogovskaya 1a, Moscow, 119435, Russia; ITMO University, Kronverksky Prospekt 49 bldg. A, Saint Petersburg, 197101, Russia; Moscow Institute of Physics and Technology, Institutsky pereulok, 9, Dolgoprudny, 141701, Moscow region, Russia

**Keywords:** gut microbiota, cancer immunotherapy, melanoma, metagenomics, fecal transplantation, compositional data analysis

## Abstract

Immunotherapy has proven to be a boon for patients grappling with metastatic melanoma, significantly enhancing their clinical condition and overall quality of life. A compelling connection was discovered between the composition of the intestinal microbiome and the effectiveness of immunotherapy substantiated in both animal models and human patients. Nonetheless, the precise biological mechanisms through which gut microbes influence melanoma treatment outcomes remain poorly understood. This study conducted a high-resolution metagenomic meta-analysis, employing cutting-edge bioinformatics techniques including genome-resolved metagenomics, strain profiling, comparative genomics, and metabolic reconstruction. According to the obtained results, the systemic metabolic depletion of the gut microbiome causes a lack of response to immunotherapy. Specifically, the presence of bacteria adept at utilizing polysaccharides, as well as those responsible for cobalamin, amino acids, and fatty acids production, decreased in patients who experienced unfavorable treatment outcomes. In contrast, patients who had successful outcomes after immunotherapy exhibited a prevalence of amino acids and cobalamin prototrophs, while autotrophy in these substances characterized the microbiomes of patients with unsuccessful outcomes. The metabolic reconstruction of short-chain fatty acid biosynthesis pathways did not differentiate bacteria linked to treatment outcomes based on their ability to produce acetate, butyrate, or propionate. However, the cobalamin-dependent Wood-Ljungdahl pathway of acetate synthesis was directly associated with immunotherapy effectiveness.

## 1 Introduction

Cutaneous melanoma stands out as one of the most formidable primary skin neoplasms, and its incidence has been on a global upswing in recent decades. However, the emergence of novel immunotherapeutic approaches and tailored medications has significantly increased patient survival as well as their quality of life [1]. Significant advances in the treatment of melanoma and other cancers have been made using immune checkpoint inhibitors (ICTs), which target proteins such as cytotoxic Tlymphocyte-associated antigen 4 (CTLA-4) and programmed cell death protein 1 (PD-1). The efficacy of ICT-based therapy has been sensational, leading to durable remission in upwards of 50% of patients grappling with metastatic melanoma [2]. However, it is imperative to acknowledge that for the remaining half of these patients, ICT therapy proves ineffectual. What’s more, in some instances, the treatment engenders undesirable side effects, ranging from dermatitis to colitis, hepatitis, antibody-related thyroid dysfunction, and pneumonia [3–6]. In the context of these studies, researchers are searching for specific host or tumor traits that may serve as predictive indicators of a patient’s response to ICT therapy and, ultimately, enhance the outcomes of immunotherapy.

The extent of impact of the human intestinal microbiota on the immunotherapy effectiveness for malignant tumors has become a focal point of extensive research within the international scientific community. The first steps into this area of study began with experiments on model animals, providing compelling evidence that intestinal bacteria may play a pivotal role in shaping the favorable outcomes of different types of anticancer therapy including ICT and promotes antitumor immunity [7–10]. These findings were subsequently validated through studies conducted on melanoma patients undergoing immunotherapy [11–13]. It is noteworthy that the composition of microbiota not only influences the effectiveness of immunotherapy but also impacts the frequency of associated side effects [14]. These pivotal discoveries were further substantiated through procedures involving fecal microbiota transplantation into gnotobiotic mice [15–17], clinical patients [18, 19] and healthy donors [20]. The results outlined above have not only showcased how specific attributes of the human gut microbiota exert influence over the outcomes of immunotherapy but have also opened up the intriguing possibility of their transferability. The phenomenon of the R (responder) phenotype being transmitted through ”fecal matter” hints at the potential involvement of particular bacterial species, a combination of bacterial species, or bacterial derivatives that could be isolated and harnessed as adjuvants to amplify the efficacy of immunotherapeutic treatments. However, despite the abundance of published research and meta-analyses on this topic, the global scientific community still grapples with an incomplete understanding of the intricate biological mechanisms underlying the regulation of the immune system by intestinal microbes in the context of cancer immunotherapy [21, 22].

In a recent meta-analysis, we made significant strides by identifying reliable gut microbiota biomarkers associated with the success of melanoma immunotherapy [23]. Based on the findings from our bioinformatics analysis, three bacteria — *Faecalibacterium prausnitzii*, *Eubacterium rectale*, and *Bifidobacterium adolescentis* — emerged as reproducible predictors of a positive response to immunotherapy. Regrettably, we did not identify any taxonomic predictors of a negative response to immunotherapy. To delve deeper into intricate biological mechanisms described above, we have expanded upon the concepts introduced in our previous research. Our current study employs a high-resolution meta-analysis, incorporating advanced bioinformatics techniques such as genome-resolved metagenomics, strain profiling, comparative genomics, and metabolic reconstruction. We have also integrated methods based on compositional data analysis (CoDa) in an original data analysis pipeline. Together, these endeavors aim to unravel the biological mechanisms underlying the gut microbiota’s influence on the regulation of antitumor immunity.

## Results

### Assembly of non-redundant MAGs catalog using melanoma patients metagenomes

In our research, we embarked on a comprehensive study of the gut microbiota in individuals battling metastatic melanoma. To accomplish this, we collected publicly available metagenomic data from recently published papers. This study focused on stool metagenomes obtained from melanoma patients collected prior to ICT administration, encompassing various treatment outcomes gleaned from five prior studies [12, 15, 16, 22, 24]. Additionally, we incorporated data showcasing the positive impact of fecal transplantation (FMT) on the results of immunotherapeutic treatment [18, 19]. In summary, stool metagenomes from a total of 680 individual samples (including 374 responders (R) and 306 nonresponders (NR)) from 7 studies were included in the analysis. A comprehensive overview of the general characteristics of this dataset is available in our previously published article [23].

Armed with this data, we proceeded to construct Metagenome-Assembled Genomes (MAGs) for each individual sample, following the protocol that we described in detail in the Materials and methods section. This endeavor yielded a total of 12,449 MAGs, subsequently compiled and dereplicated at 98% of the average nucleotide identity. The final set comprised 1,422 non-redundant MAGs, with following quality metrics: completeness 93.3 *±* 6.4 and contamination 1.6 *±* 2.0. Additional metrics values such as N50 and assembly length also presented in Supplementary Figure S1. Our genome catalog adheres to the quality standards established by the Genomic Standards Consortium (GSC) criteria [25], boasting 1,006 high-quality (71%) and 416 medium-quality (*∼*29%) MAGs. Taxonomic classification of the MAGs catalog was conducted utilizing the Genome Taxonomy Database (GTDB). Within this catalog, we identified a total of 1,416 bacterial genomes and 6 archaeal genomes, categorized into 13 distinct phyla. The most prevalent phyla include Firmicutes (902 MAGs, *∼*63%), Actinobacteria (261 MAGs, *∼*18%), Bacteroidetes (148 MAGs, *∼*10%), Proteobacteria (59 MAGs, *∼*4%), and other less frequent phyla (52 MAGs, *∼*4%). Overall assembly statistics and taxonomic annotation for the MAGs catalog can be found in Supplementary Table S1. Phylogenetic tree constructed using 1422 MAGs sequences presented in Figure 1A.

**Figure 1.**
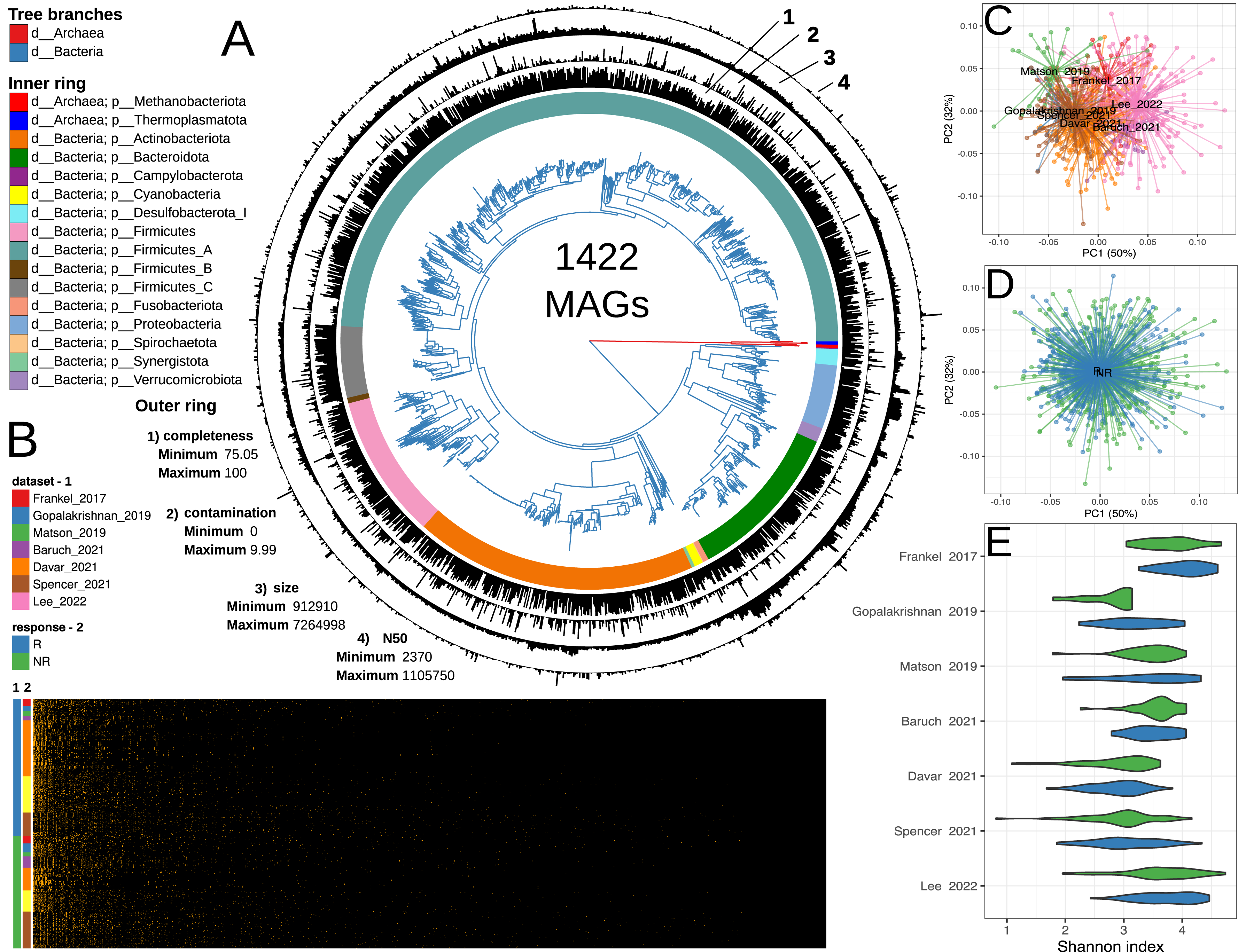
Summary of the metagenome-assembled genomes catalog assembly, taxonomic annotation, metagenomic sample mapping and metagenomic analysis. (A) Approximately maximum-likelihood phylogenetic tree generated using CheckM with 43 AA marker sequences and 1422 MAGs assembled from 680 stool metagenomes of melanoma patients. Tree branches are color-coded by bacterial or archaeal affiliation. The inner ring displays taxonomic annotations at the phylum level, aligned with the phylogenetic tree, while the outer ring presents MAGs assembly statistics. (B) Heatmap visualization illustrating the mapping results of metagenomic reads to the MAGs catalog using the InStrain tool. (C-D) Multidimensional scaling biplot showcasing MAGs-based profiles of stool metagenomes from various studies (C) and with distinct immunotherapy outcomes (D). (E) Shannon index values depicting the diversity of MAG-based profiles of stool metagenomes, stratified by dataset and immunotherapy outcome.

The next step of our analysis involved obtaining abundance profiles for the MAGs within our in the studied metagenomic catalog. This was accomplished using the InStrain approach [26]. Concurrently, we assessed alterations in routine microbial ecology metrics, including alpha- and beta-diversity. Obtained MAGs relative abundance profiles presented in Supplementary Table S2 and Figure 1B. According to the analysis of variance, the alpha-diversity statistically significantly depended on the dataset variable but not on immunotherapy outcome (FDR = 6.9e-12) while beta-diversity depended on both studied variables (FDR = 0.0001; see Supplementary Table S3 and Figure 1B-E). To identify a list of specific bacteria associated with a response to immunotherapy, we pursued a strategic approach outlined in the subsequent section.

### Discovery of consistent MAG biomarkers linked to immunotherapy outcome

Metagenomic data are compositional [Gloor et al., 2017], which imposes restrictions on the use of statistical methods directly. Several methods have been developed to represent compositional data in Cartesian space. In our study, we harnessed the capabilities of Qurro [27], a tool specifically designed to work with Songbird results [28]. Here’s an overview of our overall analysis protocol: the Songbird approach was employed to uncover microbial biomarkers linked to immunotherapy outcomes. The primary output from Songbird is a file containing differentials, which describe the log-fold change in features with respect to specific field(s) in sample metadata. The differentials are ranked, providing information on the relative associations of species with a given covariate, in our case, the immunotherapy outcome group [28]. However, it’s important to note that Songbird does not generate p-values, which can pose challenges when estimating statistical significance using Songbird alone. To overcome this limitation, we employed the Qurro method. The Qurro method calculates log ratios based on the Songbird-ranked bacterial features. These log-ratio values, being real numbers by nature, can be estimated using standard statistical methods. This approach offers convenience, as multiple bacterial features can be represented as a single value, which is beneficial for ecological modeling and statistical assessment, enabling the interpretation of results within the context of ecological ”states” without the need for separate hypothesis tests for each bacterium. Additionally, this approach enables tracking the transition of the microbiome structure from one ecological ”state” to another.

Biomarker identification was carried out following the protocol outlined in our previous publication [23]. In the first step, we identified genomes associated with R and NR by employing a differential ranking approach using Songbird individually for each data set. For subsequent analysis, we chose MAGs with an absolute Songbird differential value *>* 0.3. The logarithmic evaluation of the selected MAGs relative abundances shows a clear statistically significant difference between the R and NR groups (see Figure 2A). Evaluating the FMT data, we found that FMT responders prior to fecal transfer, exhibited statistically significantly increased Qurro log-ratio values in comparison to o FMT non-responders patients. This observation suggests the potential presence of advantageous intestine microbiome characteristics that allow these patients to respond to immunotherapy (see Figure 2B). Remarkably, this effect was observed for various donors and was reproducible in both FMT datasets (Wilcoxon rank sum test, for Baruch 2021 p = 0.008, for Davar 2021 p = 0.002). It is worth noting that according to the Baruch 2021 dataset analysis the donor variable is also important. Beneficial donors were found to have the capacity to improve the condition of recipients, effectively shifting them from an unfavorable microbiome state to a favorable one for immunotherapy. However, this transformation did not necessarily translate into individuals assuming the R phenotype. Moreover, donors with less favorable microbiomes were observed to exacerbate the condition of recipients, as depicted in Figure 2B.

**Figure 2.**
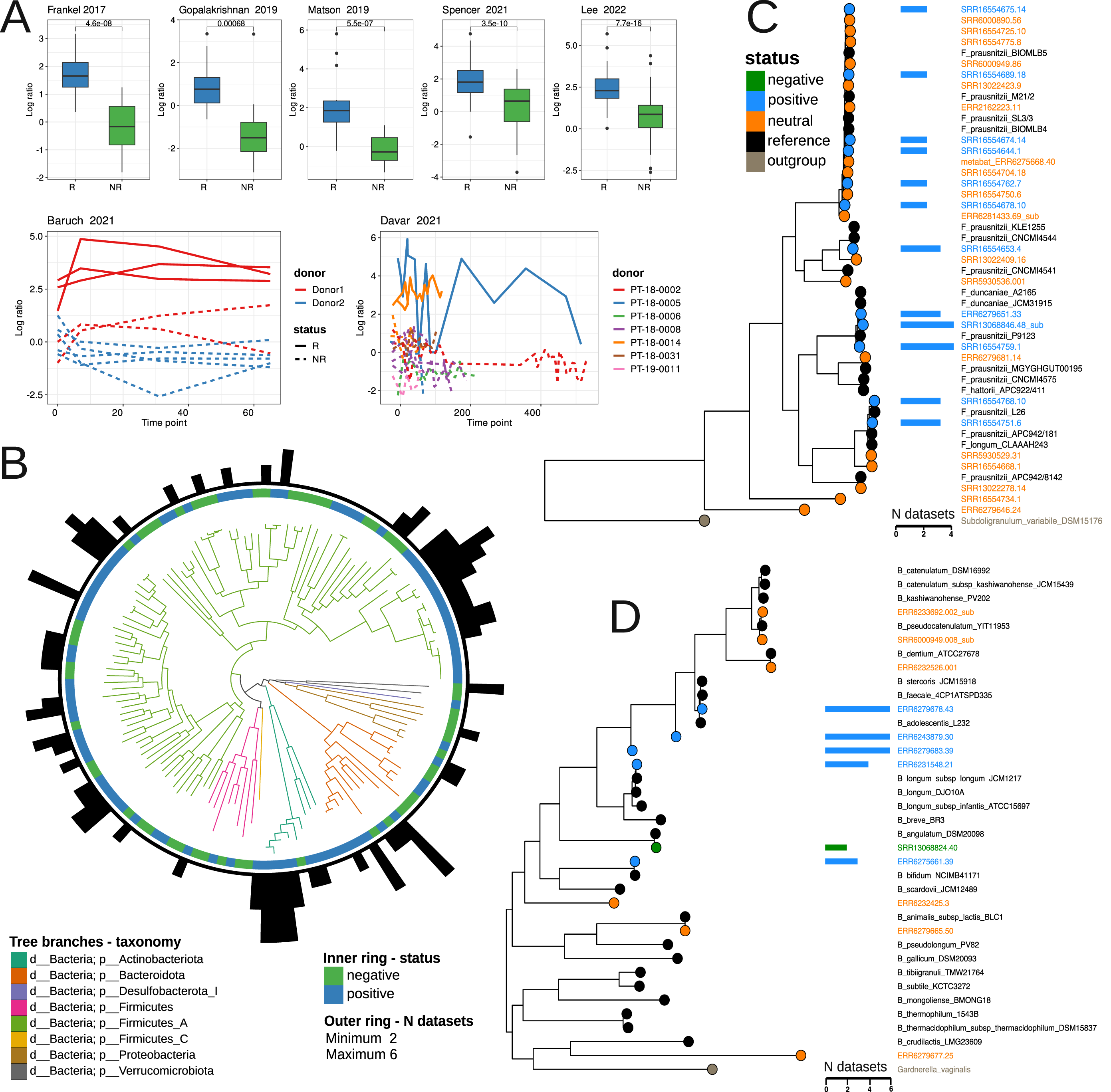
Discovery and characterization of MAG biomarkers. (A) Logarithmic ratio plots of selected features obtained through Qurro and Songbird software. MAG features with absolute differentials of 0.3 were chosen for analysis. Selected Species highlighted in blue and green correspond to Rs and NRs, respectively. Log-ratio statistical assessment via the Wilcoxon rank-sum test reveals significant differences between R and NR groups, with p-values shown. (B) Phylogenetic tree based on AA sequences of identified MAG biomarkers obtained using Orthofinder. Tree branches are colorcoded by taxonomic annotations at the phylum level. The inner ring links MAG biomarkers to the R or NR group, while the outer ring indicates the dataset numbers where the biomarker was discovered. (C-D) Orthofinder-constructed phylogenetic trees based on Faecalibacterium (C) or Bifidobacterium (D) MAGs and references. Leaf colors correspond to different biomarker sets, and additional bar plots display dataset numbers where the biomarker was found.

Following the discovery of biomarkers, we identified a total of 137 MAGs that effectively differentiated patients based on their immunotherapy outcomes. Among these, 84 MAGs were associated with positive immunotherapy outcomes, while 54 were linked to negative outcomes. These MAGs belong to 6 phyla, with their distribution as follows: Firmicutes (38 negative, 65 positive), Bacteroidetes (7 negative, 9 positive), Actinobacteria (1 negative, 8 positive), Proteobacteria (5 negative, 1 positive), Verrucomicrobiota (2 negative, 0 positive), Desulfobacterota (0 negative, 1 positive) (see Supplementary Table S4). Worth mentioning five MAGs — comprising two *Bifidobacterium adolescentis*, an unclassified Bifidobacterium, *Gemmiger quincibalis*, and *Barnesiella intestinihominis* — emerged as biomarkers associated with positive immunotherapy outcomes across all six studies. In contrast, biomarkers for negative outcomes, such as *Akkermansia* sp004167605 and *Scatavimonas* sp900540275, were replicable in no more than four datasets.

It should be noted that 30 MAGs according to the level of nucleotide similarity ¡ 98% were taxonomically annotated as *Faecalibacterium* spp. Strains of *Faecalibacterium prausnitzii* were prominent as biomarkers in six datasets. Among these, eleven *F. prausnitzii* strains were linked to positive outcomes in immunotherapy, while nineteen had a neutral status based on our biomarker discovery protocol. To gain deeper insights, we reconstructed a phylogenetic tree that encompassed all 30 Faecalibacterium MAGs, along with reference genomes of Faecalibacterium species inhabiting the human gut. As an outgroup, we incorporated the genome sequence of *Subdoligranulum variabile* DSM 15176 (see Figure 2C). All MAGs, including the 12 biomarker MAGs, were distributed among different clades within the tree. This could be attributed to the high plasticity of the genomes of *F. prausnitzii* species, suggesting that these MAGs likely belong to distinct phylogroups. Furthermore, our phylogenetic analysis indicated that three positive biomarker MAGs, namely SRR13068846.48 sub, SRR16554759.1 and ERR6279651.33, belong to the species Faecalibacterium duncaniae (strain *F. prausnitzii* P9123, which, despite its name, belongs to the *F. duncaniae* group [29]). It is worth noting that only the *F. duncaniae* clade did not contain neutral MAG biomarkers. These findings shed light on the intricate relationships within the genus Faecalibacterium in the context of immunotherapy outcomes.

The final set of non-redundant MAGs included 12 MAGs assigned to the Bifidobacterium genus. These MAGs exhibited varying associations with immunotherapy outcomes. Specifically, five of them were classified as positive MAG biomarkers (ERR6279678.43, ERR6243879.30, ERR6279683.39, ERR6231548.21, and ERR6275661.39), one (SRR13068824.40) as a negative biomarker, while the remaining six not included in the list of 137 MAG biomarkers. The taxonomic classification of these six MAGs is as follows: ERR6279678.43 and ERR6243879.30 were classified as *Bifidobacterium adolescentis*, ERR6231548.21 as *Bifidobacterium longum*, ERR6275661.39 as *Bifidobacterium bifidum*, SRR13068824.40 as *Bifidobacterium angulatum*, and ERR6279683.39 was assigned to the Bifidobacterium genus without further species annotation. We decided to look where MAG ERR6279683.39 is located on the Bifidobacterium species tree. In this regard we reconstructed a phylogenetic tree, which included all 12 MAGs of bifidobacteria and reference genomes of Bifidobacterium species inhabiting the human gut. As the outgroup, we employed the genome sequence of *Gardnerella vaginalis* UMB0386 (Figure 2D). The analysis revealed that MAG ERR6279683.39, and unexpectedly, MAG ERR6243879.30, occupied positions on the tree situated between branches related to the *B. adolescentis* group and the *B. longum* group. This observation prompted us to conduct further checks of the bifidobacteria MAGs for the chimeric assembly by GUNC. While both MAGs passed the test based on ’pass.GUNC’ in the output file, a closer examination of the output files in the ’gene calls’ and ’diamond output’ folders indicated that for MAG ERR6279683.39, 695 genes were assigned to species *B. longum*, and nearly the same number, 571 genes, were assigned to species *B. adolescentis*. Based on this, we believe that MAG ERR6279683.39 may indeed be a chimeric MAG, which likely explains its intermediary position on the tree between two species. As for MAG ERR6243879.30, there were 895 genes assigned to species *B. adolescentis* and 237 genes assigned to species *B. longum*. This indicates a potential minor contamination in this MAG, which could account for its placement outside the branch of the *B. adolescentis* group.

### Functional assessment of identified MAG biomarkers in melanoma immunotherapy outcome prediction

The annotation of the identified MAGs biomarkers involved the use of various functional databases, including CAZy, KEGG, and MetaCyc. This comprehensive annotation effort resulted in the assignment of 218 CAZy categories, 5111 KEGG orthologous groups (KOG), and 3676 MetaCyc Reactions (RXN). Derived functional profiles are available in Supplementary Tables S5-7. An analysis of variance was performed to understand the relationship between the gene categories in MAG biomarkers and the phylum and immunotherapy response variables. The results indicated that the abundance of all studied gene categories in MAG biomarkers was linked to both phylum and immunotherapy response variables, as detailed in Supplementary Table S8. Specifically, the content of KOG and RXN were significantly linked to phylum and response variables while CAZy categories profiles were also linked to phylum but the relationship with the response variable was at a lower significance level.

Further statistical testing revealed that the overall abundance of KEGG and MetaCyc gene groups annotations was increased in positive biomarkers, whereas CAZy categories showed no significant changes in either of the MAG biomarkers groups. However, a noteworthy observation was made regarding Bacteroidetes MAGs, which tended to increase the number of CAZy categories in the R group (Wilcoxon rank sum test, FDR p = 0.07). In addition, the R group was enriched by glycoside hydrolase (GH) families in comparison with NR (Wilcoxon rank sum test, p = 0.006). Specifically, only positive biomarkers Bacteroides ovatus (N CAZy = 123; GH = 70), Bacteroides xylanisolvens (N CAZy = 109; GH = 64), Bacteroides uniformis (N CAZy = 92; GH = 56), Bacteroides nordii (N CAZy = 74; GH = 41), and Parabacteroides distasonis (N CAZy = 73; GH = 45) were among the top five MAGs with the highest number of CAZy categories and GH families. The complete list of Bacteroidetes MAGs containing CAZy categories is available in Supplementary Table S9. Moreover, when examining the gene count at the phylum level, it was found that only Firmicutes and Bacteroidetes exhibited a statistically significant difference across numbers of KEGG and MetaCyc gene groups, as depicted in Supplementary Figure S3. A bi-dimensional visualization based on non-metric multidimensional scaling of functional profiles is presented in Supplementary Figure S4.

Utilizing Fisher’s exact test and applying false discovery rate (FDR) multiple testing corrections, we successfully identified specific gene groups that distinguish functional categories among the MAG biomarkers. Specifically, we found 41 KOG and 63 RXN categories that exhibited significant differences (see Supplementary Table S10). Notably, all these identified gene groups were increased in the positive MAG biomarkers group. To gain an understanding of the metabolic pathways that set apart various MAGs biomarker groups, we employed Gene Set Enrichment Analysis (GSEA). The results of this analysis revealed that 7 KEGG modules, 6 KEGG pathways, and 4 MetaCyc pathways associated with amino acids (AA) and cobalamin biosynthesis were significantly increased in the R biomarkers group. It’s worth highlighting that, according to the MetaCyc-based GSEA analysis, the PWYG-321 Mycolate biosynthesis pathway appeared to be increased in the R group. Interestingly, mycolate is an exclusive component of the cell wall of mycobacteria including the pathogen Mycobacterium tuberculosis. We further investigated this finding and translated the PWYG-321-related reactions (RXNs) into Enzyme Commission (EC) nomenclature, followed by mapping to the KEGG database. This analysis revealed that the resulting ECs were associated with the ko00061: Fatty acid biosynthetic pathway (see Supplementary Figure S5). Thus, we considered the initial observation related to the Mycolate biosynthesis pathway as an artifact of the analysis. Additionally, we explored the relationship between MAG biomarkers and the aforementioned immunotherapy-beneficial metabolic pathways. The results of this analysis, presented in Figure 3B, highlighted the top 5 genera that contained the highest number of gene groups from these metabolic pathways. These genera were Faecalibacterium, Blautia, Bacteroides, Bifidobacterium, and Ruminococcus.

**Figure 3.**
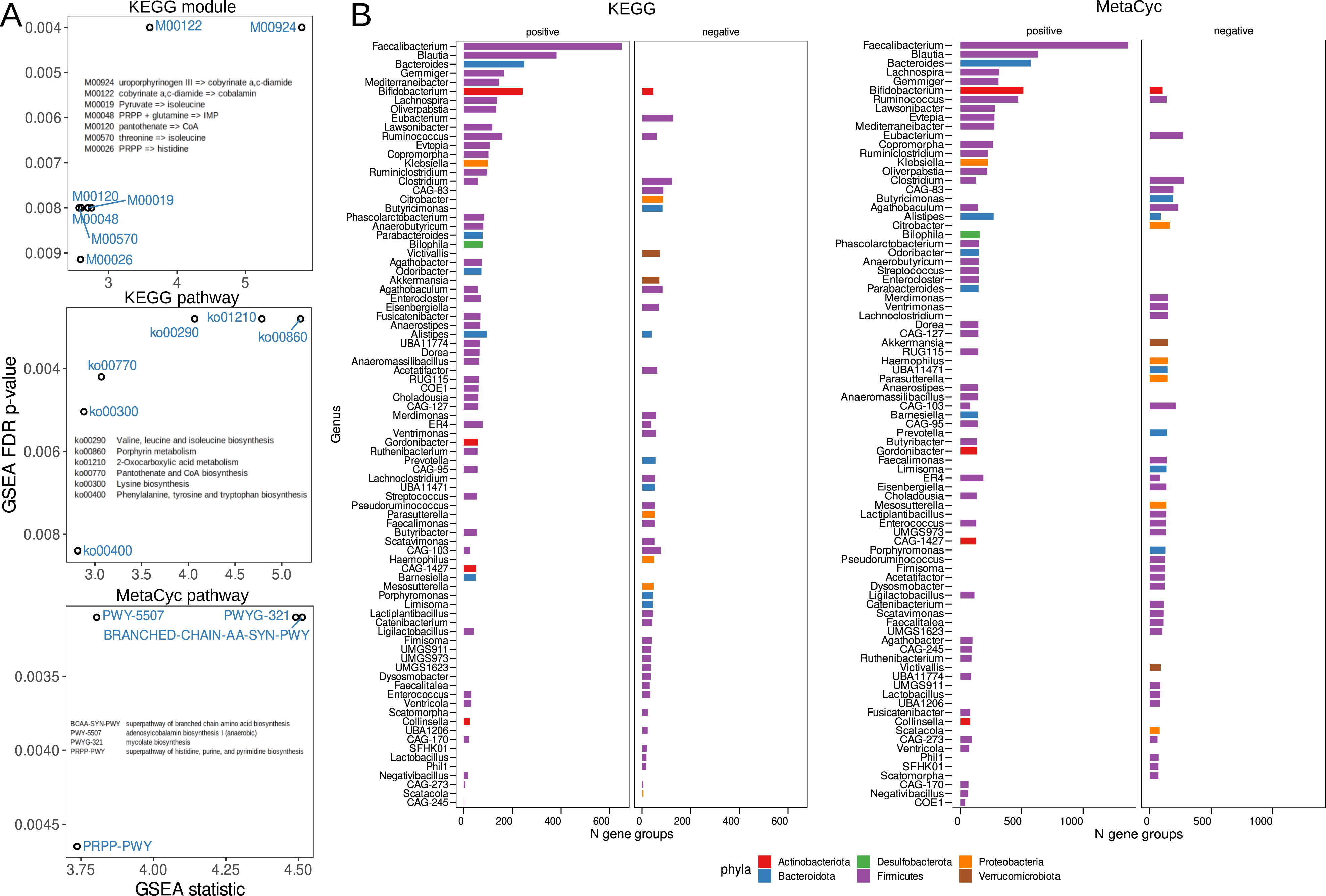
Functional differences between MAG biomarker groups (A) Gene Set Analysis (GSEA) results, with the x-axis showing GSEA analysis statistics and the y-axis representing FDR-adjusted p-values for identified functional categories. (B) Bacterial genera containing genes linked to differentially abundant KEGG and Meta-Cyc functional pathways. The x-axis indicates the total number of defined gene groups, while the y-axis corresponds to bacterial genera. Genera affiliations with bacterial phyla are color-highlighted.

### Amino Acid and cobamide auxotroph/prototroph balance linked to immunotherapy outcome

From our findings, it has become evident that the metabolic pathways related to the biosynthesis AAs and cobalamin exhibit consistent GSEA results across various gene sets. However, what piques our interest is the exploration of the contribution of both producers (prototrophs) and consumers (auxotrophs) of these vital compounds to melanoma immunotherapy outcome. Our research, employing gapseq and flux balance analysis, has successfully identified AA prototrophs and auxotrophs. The roster of target AAs includes L-Arginine, L-Asparagine, L-Cysteine, L-Glutamine, L-Histidine, L-Isoleucine, L-Leucine, L-Lysine, L-Methionine, L-Phenylalanine, L-Proline, L-Serine, L-Threonine, L-Tryptophan, L-Tyrosine, and L-Valine. Notably, our PERMANOVA results have established a statistically significant link between the distribution of AA prototrophy and auxotrophy events and both phylum and response variables (see Supplementary Table S11). Further statistical analysis has revealed that the frequency of AA prototrophy events is higher in positive immunotherapy outcome, while AA auxotrophy events are more prevalent in negative (Supplementary Figure S6A). Through Fisher exact testing, we have pinpointed specific auxotrophy/prototrophy events linked to different MAGs-biomarkers groups. Our findings indicate that L-Proline prototrophy is significantly increased only in positive biomarkers, whereas L-Tryptophan, L-Leucine, and L-Isoleucine auxotrophy are notably elevated in negative biomarkers. At lower significance levels, this trend persists, particularly with an absence of increased AA auxotrophy in positive biomarkers and no increase in prototrophy in negative biomarkers (see Supplementary Table S11). A visual representation of the distribution of AA auxotrophic/prototrophic events is presented in Figure 4A.

**Figure 4.**
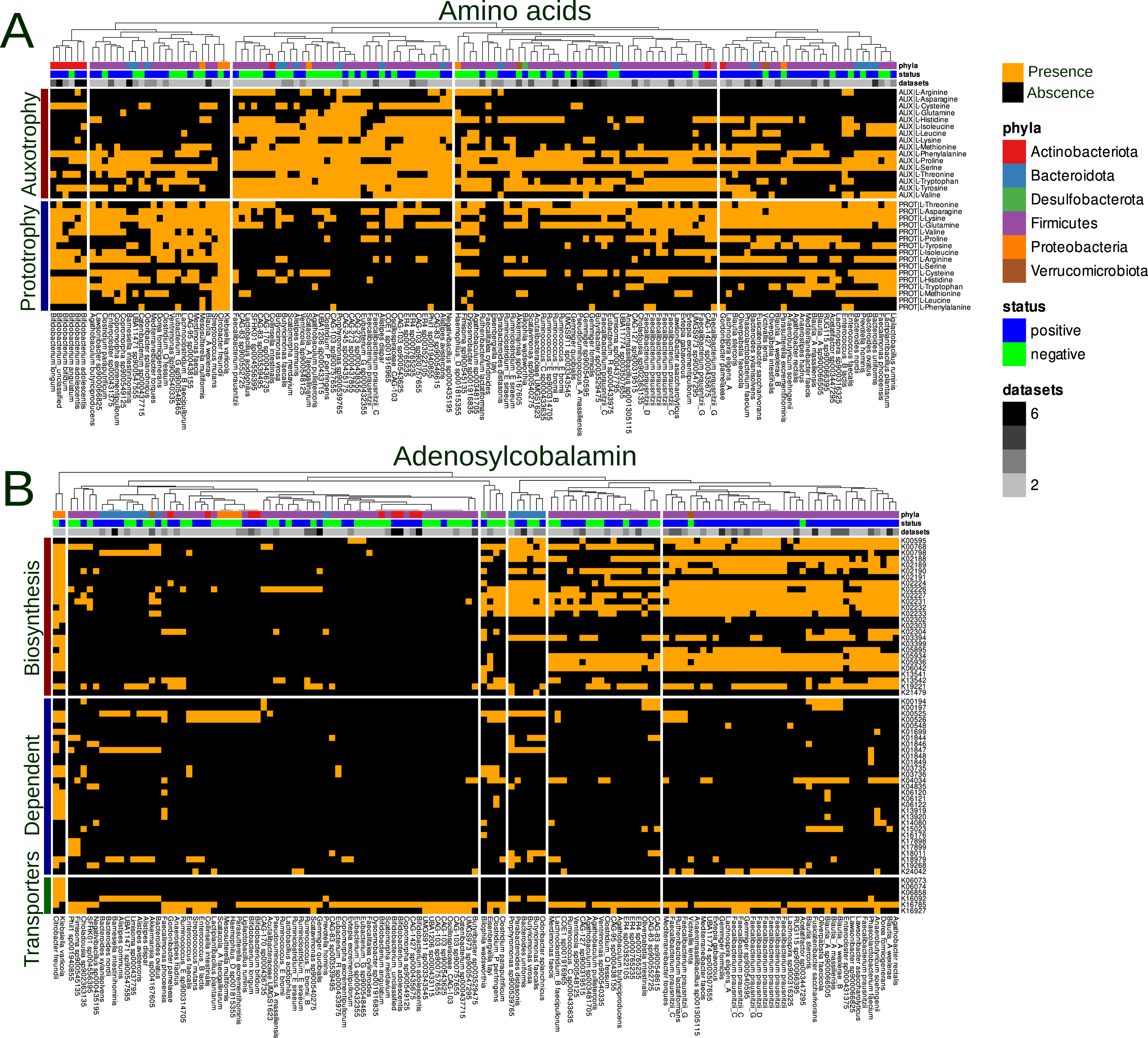
Distribution of AA auxotrophs/prototrophs and cobalamin biosynthesis genes in MAG biomarkers. (A) Bar plot displaying the distribution of predicted auxotrophy/prototrophy for specific AAs or the number of gene groups involved in cobalamin biosynthesis. The bacterial genera are defined on the x-axis. (B) Distribution of cobalamin biosynthesis genes across MAG biomarkers, with the bacterial genera plotted on the x-axis.

**Figure 5.**
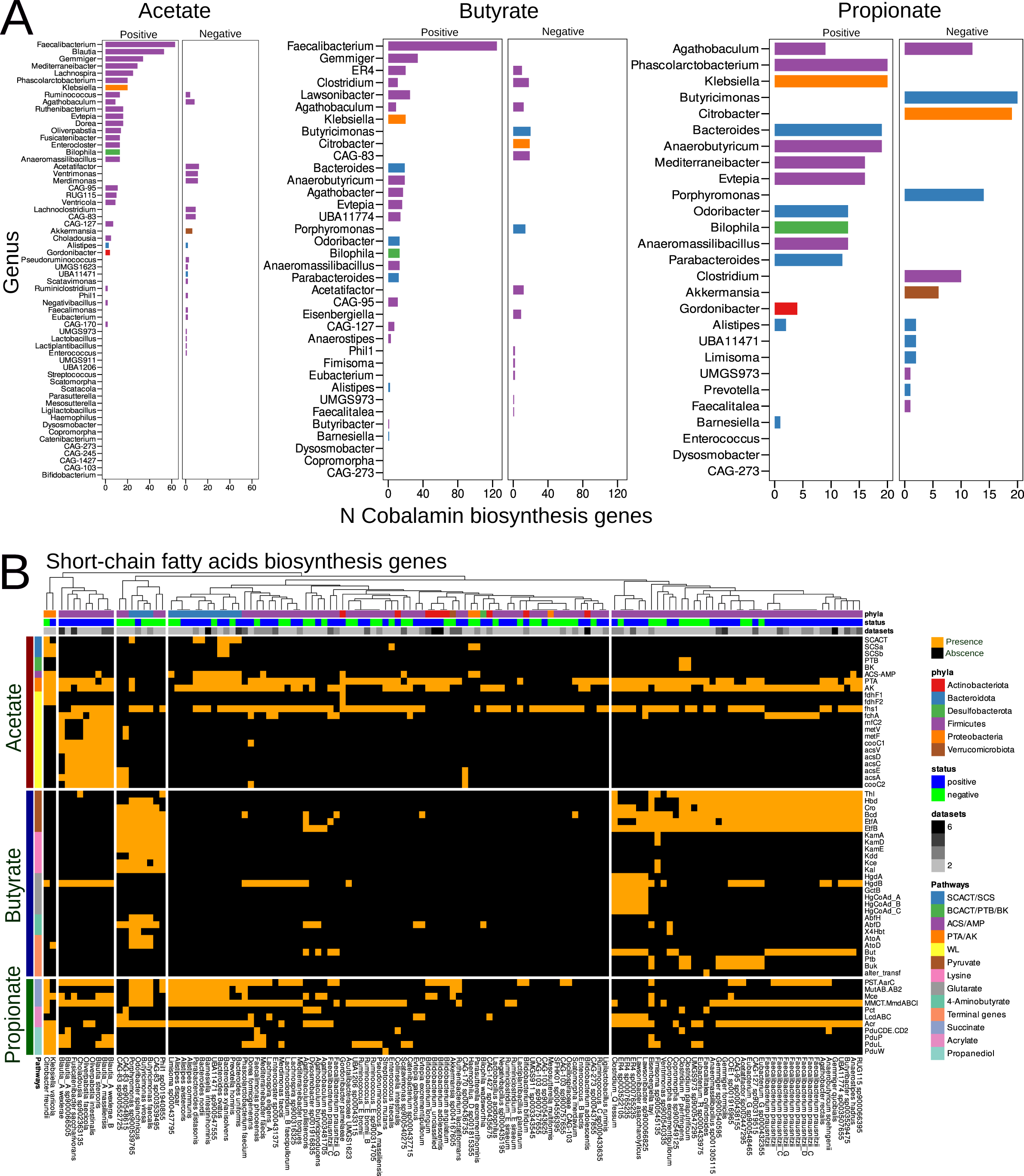
(A) Distribution of cobalamin biosynthesis genes in predicted SCFA-producers. The y-axis represents the predicted auxotrophy/prototrophy for specific AAs or the number of gene groups involved in cobalamin biosynthesis, while the bacterial genera are plotted on the x-axis. (B) Distribution of SCFA biosynthesis genes across MAG biomarkers, stratified by specific pathways. The x-axis specifies bacterial genera, and the y-axis displays genes involved in acetate, butyrate, or propionate biosynthesis.

Since gapseq did not identify the adenosylcobalamin biosynthesis pathways within the biomarker MAG sets, KOG belonging to adenosylcobalamin biosynthesis KEGG modules (M00924, M00122) were selected for further analysis. In addition, we included gene groups responsible for encoding cobalamin-dependent enzymes (N = 40) and transporters (N = 6) in our analysis. According to analysis of variance, the distribution of cobalamin-related genes was subsequently linked to both phylum and immunotherapy response variables (refer to Supplementary Table S12). Further statistical analysis revealed an increase in the number of cobalamin biosynthesis genes in positive biomarkers, while the number of genes encoding cobalamin-dependent enzymes remained unchanged between the different MAG biomarker groups (see Supplementary Figure S6B). Fisher’s exact test results indicated an increase in the occurrence of 15 cobalamin biosynthesis genes and 1 cobalamin-dependent enzyme gene in positive MAG biomarkers (see Supplementary Table S12). Upon mapping to the KEGG database, we identified KOGs linked to both the M00924 and M00122 cobalamin biosynthesis modules (see Supplementary Figure S7). A visual representation of the distribution of cobalamin-related genes among MAG biomarkers is presented in Figure 4B.

### Differential abundance of Short-Chain Fatty Acid biosynthesis pathways and their association with immunotherapy outcome

It is well-established that the SCFAs enhance overall immunity [30–32] and improve the outcomes of immunotherapy [33, 34]. Consequently, it is sensible to investigate variations in the content of SCFA biosynthetic pathways among previously identified MAG biomarkers. Our initial assesting involved the prediction of SCFA autotrophy/prototrophy using the gapseq and flux balance analysis methods. An analysis of variance using PERMANOVA indicated that SCFA production is correlated with phyla but not with immunotherapy response variables (refer to Supplementary Table S13). Furthermore, Fisher’s exact test did not reveal any statistically significant differences in the predicted producers of acetate, butyrate, and propionate among MAG groups.

Notably, certain metabolic pathways responsible for acetate and propionate production require cobalamin. Therefore, it is valuable to examine the distribution of cobalamin-producing genes among SCFA producers from the MAG biomarkers list. Our findings indicate an increase in the number of cobalamin biosynthesis genes among the positive biomarkers of acetate and butyrate producers compared to the negative biomarkers (see Supplementary Figure S8). However, a statistically significant difference in the number of cobalamin biosynthesis gene groups among predicted propionate products between the MAG groups was not observed. The distribution of cobalamin biosynthesis genes across SCFA producers is visually presented in Figure 5A.

The previous analysis results have been complemented by the catalog of predicted SCFA biosynthesis gene groups. We have identified a total of 13 SCFA pathways, including 5 related to acetate biosynthesis, 5 to butyrate biosynthesis, and 3 to propionate biosynthesis. The presence of genes included in SCFA biosynthesis pathways among MAG biomarkers is depicted in Figure 5B. According to an analysis of variance using permutations, the content of SCFA biosynthesis genes is statistically significantly associated with both phyla and immunotherapy response variables (see Supplementary Table 14). Additionally, Fisher’s exact test and further GSEA analysis revealed a linkage between the WL pathway of acetate biosynthesis and MAG biomarkers of positive immunotherapy outcomes, while butyrate biosynthesis from the lysine pathway was associated with negative immunotherapy outcomes.

## Discussion

Understanding the biological mechanisms underlying the interactions between the immune system and the human gut microbiome stands as a pivotal point in enhancing the effectiveness of cancer immunotherapy. In contrast to other cancers, sufficient quantities of the metagenomic data of patients with melanoma receiving immunotherapy have been gathered and are accessible in biological databases. By applying advanced bioinformatics techniques and reanalyzing data from these numerous research studies, we can gain deeper insights into the impact of gut bacteria on the regulation of antitumor immunity. Moreover such research endeavors will broaden our understanding of the microbiota fundamental role of its contribution in human health. In our research, we employed genome-resolved metagenomics with strain profiling, CoDa-based methods [35], and stool metagenomes from 7 previously published studies to identify sets of microbes associated with the success of immunotherapy. The list of 137 MAGs that distinguished patients based on the success of their immunotherapy were identified, according to the results of the analysis. Among the most consistently reproducible MAG biomarkers linked to positive immunotherapy outcomes were *Bifidobacterium adolescentis*, unclassified Bifidobacterium, *Barnesiella intestinihominis*, and *Gemmiger qucibalis*. It should be noted that several bifidobacteria, including *B. longum* and *B. bifidum*, were also identified as markers of successful immunotherapy outcomes, albeit in fewer datasets. Existing scientific literature corroborates these findings, with numerous published studies reporting Bifidobacterium species as indicators of favorable immunotherapy outcomes [22, 23, 36]. Additionally, results based on laboratory animals support these observations [10, 37, 38]. In contrast, a meta-analysis conducted by Limeta et al. [Limeta et al., 2020] was the sole study to establish an association between *Barnesiella intestinihominis* and improved immunotherapy outcomes. Interestingly, this bacterium has also been shown to enhance the effects of chemotherapy [39] and vascular endothelial growth factor-tyrosine kinase inhibitors treatment [40]. In turn, *G. qucibalis* has been previously identified as beneficial for the positive immunotherapy outcome of hepatobiliary cancers [41].

*F. prausnitzii* strains have also been identified as biomarkers of positive immunotherapy outcomes. According to large amount studies and meta-analyses, *F. prausnitzii* stimulates the immune system and improves response to immunotherapy of different cancer types [11, 12, 15, 16, 21, 23, 24, 41, 42]. In the intestine, *F. prausnitzii* is one of the major producers of the short-chain fatty acid (SCFA), including butyrate. Butyrate, a product of gut bacteria, enhances cytotoxic immunity and maximizes the outcomes of immunotherapy, as demonstrated by various studies [33, 34, 43, 44]. It is known that co-cultivating *F. prausnitzii* with bifidobacteria to increase colonization and boosts butyrate synthesis, likely because bifidobacteria can produce acetate, which *F. prausnitzii* requires for growth [45, 46].

Investigating the functional diversity of reproducible MAGs biomarker sets may provide understanding of processes involving the microbiota that affect antitumor immunity. Additionally, the implementation of genome-resolved metagenomics techniques allows for the examination of functions linked to specific genomes, enhancing the quality of analysis and facilitating interpretation. For example, we evaluated MAG biomarkers using the Carbohydrate-Active enZYmes Database (CAZy). High CAZy category counts in Bacteroidetes MAGs have been associated with successful immunotherapy outcomes. Among the positive biomarkers, *Bacteroides ovatus*, *Bacteroides xylanisolvens*, *Bacteroides uniformis*, *Bacteroides nordii*, and *Parabacteroides distasonis* ranked in the top 5 MAGs with the highest number of CAZy categories and GH families. *Bacteroidetes* are renowned for their ability to break down glycans using thousands of different enzyme combinations [47]. Experiments on laboratory animals demonstrated the improving effect of immunotherapy results using dietary fibers [24]. The utilization of polysaccharides by Bacteroides, namely *B. uniformis*, has been shown to influence community dynamics and butyrate synthesis in another study [48]. This suggests that glycan digestion, through cross-feeding and increased butyrate synthesis (whose immunomodulatory qualities were outlined above), promotes alterations in the microbiota that enhance the response to immunotherapy.

Further in-depth functional analysis revealed the group with positive biomarkers exhibited elevated levels of 7 KEGG modules, 6 KEGG pathways, and 4 MetaCyc pathways related to the production of compounds necessary for immunity, including amino acids, medium- and long-chain fatty acids, and cobalamin. Moreover, the list of the top five genera containing the most genes from these metabolic pathways included Faecalibacterium, Blautia, Bacteroides, Bifidobacterium, and Ruminococcus. The impact of amino acids (AAs) on supporting immune function has been extensively documented in various studies and requires no further detailed evidence or interpretation [Kelly et al., 2020]. Microbiome-produced medium- and long-chain fatty acids have the potential to stimulate antitumor immunity through binding to free fatty acid receptors. However, the connections between gut microbe-produced cobalamin and human immunity are less straightforward. Microbial-derived cobalamin and other corrinoids may serve an ecological role, be distributed by producers within the community, and be utilized by cobalamin-auxotrophic microbes [49]. On the other hand, corrinoids can be shared between microbes and Caco epithelial cells via vesicular transport [50]. Therefore, adding additional cobalamin from the gut microbes as a supplement to the diet-derived form may help the immune system’s ability to fight tumors. In addition to functional annotation analysis, the metabolic reconstruction approach revealed that AA and cobalamin prototrophs were associated with positive immunotherapy outcomes, while auxotrophs were linked to unfavorable outcomes. It is conceivable that ”altruistic” bacterial behavior may make them more beneficial to the host and build community resilience.

It is well-established that the SCFAs enhance overall immunity [30–32] and influence immunotherapy outcomes [33, 34]. MAG biomarker sets did not exhibit significant differences in their predicted capacity to produce acetate, butyrate, or propionate. However, the WL acetate production pathway was connected with successful treatment outcomes, while the butyrate biosynthesis pathway through lysine degradation was linked to unsuccessful immunotherapy, according to analysis of the reconstructed SCFA biosynthesis pathways. Notably, three bacterial genera, known for producing major SCFAs via the WL pathway (Blautia, Fusicatenibacter, and Oliverpabstia), are also involved in cobalamin biosynthesis (see Figure 4). The change in activity of the WL pathway might be driven by a cross-feeding relationship with Bifidobacterium species (for example *B. bifidum*), as they are specialized carbohydrate-fermenting species that produces the substrates for the CO_2_ fixation by the WL pathway [51]. Moreover, the WL pathway demonstrates a kind of positive feedback - providing additional acetate production, which in turn affects the increase in butyrate production [52]. This strategy appears to be more advantageous because it does not utilize other important metabolites, in this case AA, for the synthesis of SCFA. Furthermore, acetate produced by the gut microbiota can directly enhance immunological function [53, 54].

Recently published studies showed the effectiveness of fecal microbiota transplantation (FMT) received from R patients [18, 19] or healthy volunteers for improving immunotherapy outcome [20]. According to our results, the microbiota structure of the R patients differed from the microbiota of NRs already before the performing of FMT procedure. It is therefore reasonable to suppose that the R patient’s microbiota responded to the donor’s feces in a way that improved antitumor immunity while NR patient’s could not respond properly. Perhaps, due to the absence (inadequate amount) of specific bacteria in the initial NRs’ microbiota, donor feces were unable to result in such an improvement. Previously, in experiments on laboratory animals, the possibility of replenishing cobalamin deficiency by eating feces was noted [55, 56]. It can be assumed that absorption of donor feces-derived cobalamin enhances cytotoxic immunity which may have a beneficial effect on immunotherapy results. Moreover, fecal-derived corrinoids inaccessible to humans can be used by the corrinoids auxotrophs in the intestine for improved growth and metabolism. Since the specter of colonizers was similar between R and NR patients in our previous analysis, it’s possible that the FMT mechanism in this particular case is linked to fecal cobalamin (or other metabolites) to a greater extent than donor-derived microbes colonization. On the other hand, it is impossible to disregard the potential impact of donor microorganisms. The condition of the Rs’ overall gut microbiota and the effectiveness of immunotherapy may both be improved by donor microorganisms’ ability to restore lost ecological connections which degraded at Rs’ to a lesser extent compared to NRs’.

According to the results obtained and scientific literature data, it is evident that dietary fiber consumption can have a positive impact on melanoma immunotherapy outcomes [24]. Nevertheless, in our opinion, further research involving large patient cohorts is essential to identify the most effective types of fiber and develop precise administration regimens for clinical use. While studies on laboratory animals with melanoma models have shown that the use of bifidobacteria and lactobacteria can enhance antitumor immunity [10, 37, 38, 57, 58], it’s important to note that Bifidobacterium spp. are also emerging as reproducible biomarkers of positive immunotherapy outcomes in patient-based studies and meta-analyses [22, 23, 36]. Consequently, the incorporation of ’classical probiotics’ into immunotherapy regimens holds promise on the one hand. On the other hand, reported the influence of commercial probiotics on immunotherapy outcomes has yielded negative effects [24]. Bifidobacteria may serve as markers of a microbiota’s ’correct’ state in humans. This concept finds support in clinical trial results, where patients receiving products containing *Clostridium byturicum* not only exhibited increased levels of bifidobacteria in their gut but also experienced improved survival [59]. Another potential probiotic candidate for enhancing immunotherapy outcomes could be *Propionibacterium freudenreichii* due to its ability to produce cobalamin. However, the most promising prospects lie in the potential development of probiotics based on *F. prausnitzii* for clinical use [60]. As noted above, this bacterium has been linked to the enhancement of antitumor immunity in numerous studies, including our own analysis. Thus, the use of other novel types of probiotics can be considered in the future as a potential direction for improving antitumor immunity.

## Conclusion

Summarizing the findings and hypothesis, it is evident that the human gut microbiota plays a crucial role in establishing mutualistic relationships aimed at enhancing antitumor immunity. Here are key points of the conclusions drawn from the analysis: (1) polysaccharide utilization: Bacteroidetes species decompose the complex carbohydrates and produce substrates for other community members including *F. prausnitzii* ; (2) cobalamin sharing: *F. prausnitzii* produces and shares cobalamin with Bacteroidetes and other cobalamin auxotrophs. Bacteroidetes may potentially contribute to cobalamin transport within gut epithelial cells via extracellular vesicles, facilitating its distribution to other microbial community microbes and the host. This cooperative exchange ensures that essential nutrients are available to various members of the microbiota building community resilience; (3) acetate/butyrate metabolism:

*F. prausnitzii* utilizes acetate produced by *Bifidobacterium* spp. and/or through the WL pathway for its growth and butyrate production. This cross-feeding relationship results in the increased production of both acetate and butyrate, benefiting the host. The use of additional carbon sources such as CO_2_ in the WL pathway may allow the production of more acetate and butyrate and receipt of free resources for the production of other important metabolites; (4) cross-feeding: interestingly, the analysis suggests that it’s not merely the ability to produce butyrate but rather the cross-feeding between bacteria, linked to butyrate production, that contribute to improved immunotherapy outcomes. This highlights the importance of microbial interactions in enhancing immunity; (5) Altruistic behavior: Immunotherapy beneficial microbiota exhibit altruistic behavior producing important metabolites such as amino acids and cobalamin and can spread them between community members which improves the host’s anti-tumor immunity status. While the harmful microbiota behaves selfishly and competes for these important resources with the host. In summary, our analysis has uncovered intricate links between biological hypotheses and obtained results, shedding light on the complex relationships within the human gut microbiota and their impact on antitumor immunity. Our data analysis protocol can be used to study other types of cancer and pave the way for further research in this field.

## Methods

### Metagenomic datasets, analysis, and data preprocessing

Sequencing data from gut metagenome samples of melanoma patients were collected from seven published studies [12, 15, 16, 18, 19, 22, 24]. These data underwent pre-processing which was performed as follows. Metagenomic data were subjected to quality assessment using FastQC [https://github.com/s-andrews/FastQC]. Technical sequences and low-quality bases were removed from the data using the Trimmomatic tool [61]. Human sequences present in the metagenomic samples were eliminated using bbmap [62] and human genome GRCh37 [https://www.ncbi.nlm.nih.gov/genome/ guide/human]. All the pre-processing computational procedures were executed using the Assnake metagenomics pipeline [https://github.com/ASSNAKE]. Detailed information about the characteristics of the metagenomic datasets and pre-processing statistics can be found in our previous study [23].

### Construction of non-redundant metagenome-assembled genomes (MAGs) catalog

The sequencing data obtained from the pre-processing step were utilized to construct metagenomic contigs using the MEGAHIT assembler [63]. Contigs with a length greater than 1000 base pairs were retained for further analysis. Subsequently, resulting assembly results were subjected to binning using two methods: MaxBin2 [64] and MetaBAT2 [65]. To create an optimized, non-redundant bin set for each sample, the DASTool [66] was employed. For the construction of a non-redundant catalog of MAGs, the dRep tool [67] was used with specific parameters: –completeness 75 and –contamination 10. To assess the final quality of the resulting bin set, the CheckM framework [68] was applied. Taxonomic annotation of the resulting MAGs catalog was achieved using the GTDB-Tk tool [69, 70]. A phylogenetic tree incorporating all MAGs sequences was constructed utilizing the FastTree2 [71], based on the CheckM 43 marker AAs. Multiple alignment was performed using MUSCLE [72]. For visualization of the MAG biomarkers phylogenetic tree, the Empsess [73], included in the QIIME2 framework [74], was employed. Both InStrain and the MAGs catalog were used for metagenomic data profiling [26]. Using the abundance matrix of MAGs obtained in the previous computational step, the Shannon index was calculated using the diversity function within the vegan v2.6-4 package for GNU/R [https://github.com/vegandevs/vegan]. For beta-diversity assessment and two-dimensional visualization, robust principal component analysis (rPCA) [75] was employed. Links between the dataset and response variables to microbial composition were evaluated using PERMANOVA and the robust Aitchison distance (calculated by DEICODE) with 10,000 permutations, implemented in the adonis2 function of the vegan v2.6-4 package. Visualization of the obtained results was performed using the ggplot2 v3.4.2 library for GNU/R [https://ggplot2.tidyverse.org].

### Strategy for discovering MAG biomarkers

Identification of MAGs-biomarkers linked to immunotherapy outcome was performed in a similar way as in the previous article [23]. At the first step, MAGs whose relative abundance linked to immunotherapy outcome were identified using the Songbird tool [28]. The log ratios of the selected MAGs were calculated using Qurro [27] while the statistical significance of log ratios was accessed using the Wilcoxon rank-sum test implemented in the basic functional of GNU/R. The second step was the creation of a list of consistent MAGs-biomarkers by following this methodology: 1) microbial species that were associated with positive immunotherapy outcome in more than one dataset were added to the list; 2) MAGs that were associated with negative outcome in at least one dataset were excluded from the list of MAGs-biomarkers regardless of the number of datasets in which they were associated with a positive outcome. The specific MAGs biomarkers linked to negative outcomes were identified in a similar way. The ggplot2 v3.4.2 [https://ggplot2.tidyverse.org] library for GNU/R was used for visualization of obtained results. Orthofinder was used for construction of a phylogenetic tree using identified MAGs biomarkers sequences. Empsess tool was used for visualization of obtained MAGs-biomarkers phylogenetic tree.

### Phylogenetic tree construction of Faecalibacterium and Bifidobacterium Species

MAGs assigned to the genera Faecalibacterium (30 MAGs) and Bifidobacterium (12 MAGs) were taken for phylogenetic analysis. Open reading frames and translated AA sequences from selected MAGs were predicted using Prodigal version 2.6.3 [76]. The phylogenomic trees based on predicted sequences were reconstructed using OrthoFinder version 2.5.4 [77] with default parameters. Genomes from species inhabiting the human gut were selected as references. The genome sequence from Gardnerella vaginalis strain UMB0386 (GenBank: PKJK01000001.1) was used as an outgroup for Bifidobacterium, while as an outgroup for Faecalibacterium the genome sequence of Subdoligranulum variabile strain DSM 15176 (GenBank: ACBY02000001.1). The trees were visualized using the ggplot2 v3.4.2 and ggtree v3.6.2 [78] packages for GNU/R. To additional control the Bifidobacterium MAGs quality, the sequences were checked by GUNC v1.0.5 [79] to filter out chimeric genomes based on column ’pass.GUNC’ in the gunc output file.

### Functional profiling of MAG biomarkers

To explore the presence of CAZy within MAGs, we employed a series of bioinformatic analyses. Firstly, AA sequences, predicted by Prodigal v2.6.3 [76], were aligned against bacterial protein sequences from the CAZy database [http://www.cazy.org] [80] and the KEGG database [https://www.genome.jp/kegg] [81] using the blastp mode of DIAMOND algorithm v2.0.15 [82] with a stringent threshold of 80% identity and 80% query coverage. Additionally, we used the gapseq method [83] in conjunction with the MetaCyc database [https://metacyc.org] [84] for further functional annotation of MAG biomarkers.

For bi-dimensional visualization of functional data, we employed non-metric multidimensional scaling and the Bray-Curtis dissimilarity metric [85]. To measure distances and dissimilarities in functional content comparisons among MAG biomarkers, we conducted PERMANOVA using the adonis2 function from the vegan package. Differences in the abundance of identified functional categories between MAG biomarker sets were assessed using the Wilcoxon rank sum test with FDR correction for multiple testing. Furthermore, differences in functional content among MAGs groups were determined using Fisher exact tests with FDR correction, implemented in GNU/R.

To discern disparities in KEGG/MetaCyc gene sets between MAGs groups, we employed gene set analysis (GSA) from the piano Bioconductor package [86]. Specifically, we used the ’reporter feature algorithm’ with a gene set significance threshold of adj. p *<* 0.05. Fisher exact test-derived FDR-corrected p-values served as input data for GSA analysis. Genes with uncorrected p-values *>* 0.5 were classified as ’increased,’ while those with p-values *<* 0.5 were deemed ’decreased.’ Visualization of the results was accomplished using the ggplot2 v3.4.2 and pheatmap v1.0.12 [https://github.com/raivokolde/pheatmap] libraries for GNU/R.

In our MAGs analysis, we focused on identifying genes encoding pathways responsible for acetate, butyrate, and propionate production. For acetate, we considered six possible synthesis pathways, including the WL pathway and a recently discovered pathway involving succinyl-CoA:acetate CoA-transferase and succinyl-CoA synthetase [52, 87, 88]. Meanwhile, for butyrate and propionate, we explored four and three possible synthetic pathways, respectively [89, 90] (refer to Supplementary Table S15 for details). We assembled a reference catalog of gene products for each pathway, resulting in 4563 AA sequences for acetate pathways, 2744 for butyrate pathways, and 415 for propionate pathways. Subsequently, we conducted DIAMOND v2.0.15 blastp searches [82] and used the program gapseq v1.1 [83] with default parameters to validate the presence of these pathways. Additionally, we employed gapseq profiles and flux balance analysis to predict AAs and SCFA consumers-producers [83, 91].

## Supplementary information

### 1.1 Supplementary fugures

Figure S1. MAGs catalog assembly statistics.

Figure S2. Amount of different functional categories across MAGs and biomarker groups. Additionally, the graphs show p-values obtained using Wilcoxon rank-sum test with FDR multiple testing correction.

Figure S3. Functional category diversity across MAG biomarkers groups stratified by phyla. Additionally, the graphs show p-values obtained using Wilcoxon rank-sum test with FDR multiple testing correction.

Figure S4. Nonmetric multidimensional scaling biplots.

Figure S5. Mapping identified gene groups of MetaCyc PWYG-321 Mycolate biosynthesis to the KEGG ko00061 fatty acid biosynthesis pathway.

Figure S6. Amount of prototrophs and auxotrophs across MAG biomarker groups. Additionally, the graphs show p-values obtained using Wilcoxon rank-sum test with FDR multiple testing correction.

Figure S7. Upregulated cobalamin synthesis genes in biomarker-positive group mapped to KEGG ko00860: porphyrin Metabolism.

Figure S8. Amount of cobalamin biosynthesis among SCFA producer MAGs and biomarkers. Additionally, the graphs show p-values obtained using Wilcoxon rank-sum test with FDR multiple testing correction.

### 1.2 Supplementary tables

Table S1. Assembly statistics and taxonomic annotation of MAGs.

Table S2. Relative abundance of MAGs across melanoma patient metagenomes obtained using the InStrain method (Percentage of overall abundance).

Table S3. Statistical evaluation of alpha- and beta-diversity of taxonomic profiles obtained using the MAGs catalog.

Table S4. MAG biomarkers associated with immunotherapy outcome. Table S5. Distribution of CAZy categories across MAG biomarkers.

Table S6. Distribution of KOGs across MAG biomarkers. Table S7. Distribution of RXN across MAG biomarkers.

Table S8. Statistical assessment of gene group content variations in MAG biomarkers.

Table S9. Quantity of CAZy Categories in MAG biomarkers.

Table S10. Statistical analysis results of gene group content across MAG biomarker groups.

Table S11. Statistical assessment of AA auxotrophy/prototrophy predictions by Gapseq.

Table S12. Statistical analysis of cobalamin-linked gene groups across MAG biomarkers.

Table S13. Statistical assessment of gapseq-predicted SCFA production ability across MAG biomarkers.

Table S14. Statistical analysis of gene content related to SCFA biosynthesis across MAG biomarkers.

Table S15. Metabolic pathways for the production of acetate, butyrate, and propionate by gut microbes.

## List of abbreviations

AA: amino acid.
CoDa: compositional data analysis.
CAZy: Carbohydrate-Active Enzymes.
GH: glycoside hydrolase.
EC: Enzymes nomenclature.
FDR: false discovery rate.
FMT: fecal microbiota transplantation.
(GTDB): Genome Taxonomy Database.
GSA: gene set analysis.
KEGG: Kyoto encyclopedia genes and genomes.
KOG: KEGG orthologous groups.
MAG: metagenome assembled genomes.
NR: non-responder.
R: responder.
rPCA: robust principal components analysis.
RXN: reactions.
SCFA: short-chain fatty acid.
WL: Wood-Ljungdahl.

## Declarations

- Acknowledgments We would like to acknowledge the valuable computational resources provided by the FRCC PCM ”Genomics, Proteomics, Metabolomics” Center [http://rcpcm.org/?p=2806]. Additionally, our gratitude extends to Roman Yunes for his assistance in editing the English text.
- Funding The financial backing for this study was provided by the Russian Science

Foundation under grant number 22-75-10029, accessible at https://rscf.ru/project/22-75-10029/.

- Conflict of interest/Competing interests (check journal-specific guidelines for which heading to use) The authors declare that the research was conducted in the absence of any commercial or financial relationships that could be construed as a potential conflict of interest.
- Availability of data and materials In this study, we utilized openly accessible data retrieved from the NCBI-EBI Sequence Read Archives, identified by the following BioProjects accession numbers: PRJNA397906, PRJEB22893, PRJNA399742, PRJNA678737, PRJNA67286, PRJNA770295, and PRJEB43119. Comprehensive findings from our project are comprehensively detailed within the article text, along with supplementary materials. We have also made available a catalog of Metagenome-Assembled Genomes (MAGs), taxonomic annotation results, phylogenetic trees, and Empress profiles for the QIIME2 viewer. These resources have been shared via the figshare service, accessible through the following link: https://doi.org/10.6084/m9.figshare.24146913.v3.
- Authors’ contributions NVZ: Performed functional and phylogenetic analyses, wrote the manuscript, contributed to data analysis and visualization, and assisted in study design conceptualization and interpretation of obtained results. VAK: Provided bioinformatics analysis support. MDM: Contributed to manuscript preparation and interpretation of obtained results, as well as provided bioinformatics analysis support. ABI, VIU: Contributed to metagenomic data analysis and visualization, and participated in manuscript preparation. KMK: Contributed to the conceptualization of the study and manuscript preparation. EIO: Led study design and conceptualization, performed computational experiments, designed and implemented data analysis and visualization, contributed to functional and phylogenetic analysis, and wrote the manuscript.

## References

[1] Switzer, B., Puzanov, I., Skitzki, J.J., Hamad, L., Ernstoff, M.S.: Managing metastatic melanoma in 2022: a clinical review. JCO Oncology Practice 18(5), 335–351 (2022)

[2] Larkin, J., Chiarion-Sileni, V., Gonzalez, R., Grob, J.J., Cowey, C.L., Lao, C.D., Schadendorf, D., Dummer, R., Smylie, M., Rutkowski, P., et al.: Combined nivolumab and ipilimumab or monotherapy in untreated melanoma. New England journal of medicine 373(1), 23–34 (2015)

[3] Horvat, T.Z., Adel, N.G., Dang, T.-O., Momtaz, P., Postow, M.A., Callahan, M.K., Carvajal, R.D., Dickson, M.A., D’Angelo, S.P., Woo, K.M., et al.: Immune-related adverse events, need for systemic immunosuppression, and effects on survival and time to treatment failure in patients with melanoma treated with ipilimumab at memorial sloan kettering cancer center. Journal of Clinical Oncology 33(28), 3193 (2015)

[4] Wolchok, J.D., Chiarion-Sileni, V., Gonzalez, R., Rutkowski, P., Grob, J.-J., Cowey, C.L., Lao, C.D., Wagstaff, J., Schadendorf, D., Ferrucci, P.F., et al.: Overall survival with combined nivolumab and ipilimumab in advanced melanoma. New England Journal of Medicine 377(14), 1345–1356 (2017)

[5] Roy, S., Trinchieri, G.: Microbiota: a key orchestrator of cancer therapy. Nature Reviews Cancer 17(5), 271–285 (2017)

[6] Robert, C., Ribas, A., Schachter, J., Arance, A., Grob, J.-J., Mortier, L., Daud, A., Carlino, M.S., McNeil, C.M., Lotem, M., et al.: Pembrolizumab versus ipilimumab in advanced melanoma (keynote-006): post-hoc 5-year results from an open-label, multicentre, randomised, controlled, phase 3 study. The Lancet Oncology 20(9), 1239–1251 (2019)

[7] Iida, N., Dzutsev, A., Stewart, C.A., Smith, L., Bouladoux, N., Weingarten, R.A., Molina, D.A., Salcedo, R., Back, T., Cramer, S., et al.: Commensal bacteria control cancer response to therapy by modulating the tumor microenvironment. science 342(6161), 967–970 (2013)

[8] Viaud, S., Saccheri, F., Mignot, G., Yamazaki, T., Daillère, R., Hannani, D., Enot, D.P., Pfirschke, C., Engblom, C., Pittet, M.J., et al.: The intestinal microbiota modulates the anticancer immune effects of cyclophosphamide. science 342(6161), 971–976 (2013)

[9] Vétizou, M., Pitt, J.M., Daillère, R., Lepage, P., Waldschmitt, N., Flament, C., Rusakiewicz, S., Routy, B., Roberti, M.P., Duong, C.P., et al.: Anti-cancer immunotherapy by ctla-4 blockade relies on the gut microbiota. Science 350(6264), 1079–1084 (2015)

[10] Sivan, A., Corrales, L., Hubert, N., Williams, J.B., Aquino-Michaels, K., Earley, Z.M., Benyamin, F.W., Man Lei, Y., Jabri, B., Alegre, M.-L., et al.: Commensal bifidobacterium promotes antitumor immunity and facilitates anti–pd-l1 efficacy. Science 350(6264), 1084–1089 (2015)

[11] Chaput, N., Lepage, P., Coutzac, C., Soularue, E., Le Roux, K., Monot, C., Boselli, L., Routier, E., Cassard, L., Collins, M., et al.: Baseline gut microbiota predicts clinical response and colitis in metastatic melanoma patients treated with ipilimumab. Annals of Oncology 28(6), 1368–1379 (2017)

[12] Frankel, A.E., Coughlin, L.A., Kim, J., Froehlich, T.W., Xie, Y., Frenkel, E.P., Koh, A.Y.: Metagenomic shotgun sequencing and unbiased metabolomic profiling identify specific human gut microbiota and metabolites associated with immune checkpoint therapy efficacy in melanoma patients. Neoplasia 19(10), 848–855 (2017)

[13] Routy, B., Le Chatelier, E., Derosa, L., Duong, C.P., Alou, M.T., Daillère, R., Fluckiger, A., Messaoudene, M., Rauber, C., Roberti, M.P., et al.: Gut microbiome influences efficacy of pd-1–based immunotherapy against epithelial tumors. Science 359(6371), 91–97 (2018)

[14] Dubin, K., Callahan, M.K., Ren, B., Khanin, R., Viale, A., Ling, L., No, D., Gobourne, A., Littmann, E., Huttenhower, C., et al.: Intestinal microbiome analyses identify melanoma patients at risk for checkpoint-blockade-induced colitis. Nature communications 7(1), 10391 (2016)

[15] Matson, V., Fessler, J., Bao, R., Chongsuwat, T., Zha, Y., Alegre, M.-L., Luke, J.J., Gajewski, T.F.: The commensal microbiome is associated with anti–pd-1 efficacy in metastatic melanoma patients. Science 359(6371), 104–108 (2018)

[16] Gopalakrishnan, V., Spencer, C.N., Nezi, L., Reuben, A., Andrews, M., Karpinets, T., Prieto, P., Vicente, D., Hoffman, K., Wei, S.C., et al.: Gut microbiome modulates response to anti–pd-1 immunotherapy in melanoma patients. Science 359(6371), 97–103 (2018)

[17] Lam, K.C., Araya, R.E., Huang, A., Chen, Q., Di Modica, M., Rodrigues, R.R., Lopés, A., Johnson, S.B., Schwarz, B., Bohrnsen, E., et al.: Microbiota triggers sting-type i ifn-dependent monocyte reprogramming of the tumor microenvironment. Cell 184(21), 5338–5356 (2021)

[18] Davar, D., Dzutsev, A.K., McCulloch, J.A., Rodrigues, R.R., Chauvin, J.-M., Morrison, R.M., Deblasio, R.N., Menna, C., Ding, Q., Pagliano, O., et al.: Fecal microbiota transplant overcomes resistance to anti–pd-1 therapy in melanoma patients. Science 371(6529), 595–602 (2021)

[19] Baruch, E.N., Youngster, I., Ben-Betzalel, G., Ortenberg, R., Lahat, A., Katz, L., Adler, K., Dick-Necula, D., Raskin, S., Bloch, N., et al.: Fecal microbiota transplant promotes response in immunotherapy-refractory melanoma patients. Science 371(6529), 602–609 (2021)

[20] Routy, B., Lenehan, J.G., Miller Jr, W.H., Jamal, R., Messaoudene, M., Daisley, B.A., Hes, C., Al, K.F., Martinez-Gili, L., Punčochář, M., et al.: Fecal microbiota transplantation plus anti-pd-1 immunotherapy in advanced melanoma: A phase i trial. Nature medicine 29(8), 2121–2132 (2023)

[21] Limeta, A., Ji, B., Levin, M., Gatto, F., Nielsen, J.: Meta-analysis of the gut microbiota in predicting response to cancer immunotherapy in metastatic melanoma. JCI insight 5(23) (2020)

[22] Lee, K.A., Thomas, A.M., Bolte, L.A., Björk, J.R., Ruijter, L.K., Armanini, F., Asnicar, F., Blanco-Miguez, A., Board, R., Calbet-Llopart, N., et al.: Cross-cohort gut microbiome associations with immune checkpoint inhibitor response in advanced melanoma. Nature Medicine 28(3), 535–544 (2022)

[23] Olekhnovich, E.I., Ivanov, A.B., Babkina, A.A., Sokolov, A.A., Ulyantsev, V.I., Fedorov, D.E., Ilina, E.N.: Consistent stool metagenomic biomarkers associated with the response to melanoma immunotherapy. Msystems 8(2), 01023–22 (2023)

[24] Spencer, C.N., McQuade, J.L., Gopalakrishnan, V., McCulloch, J.A., Vetizou, M., Cogdill, A.P., Khan, M.A.W., Zhang, X., White, M.G., Peterson, C.B., et al.: Dietary fiber and probiotics influence the gut microbiome and melanoma immunotherapy response. Science 374(6575), 1632–1640 (2021)

[25] Bowers, R.M., Kyrpides, N.C., Stepanauskas, R., Harmon-Smith, M., Doud, D., Reddy, T., Schulz, F., Jarett, J., Rivers, A.R., Eloe-Fadrosh, E.A., et al.: Minimum information about a single amplified genome (misag) and a metagenome-assembled genome (mimag) of bacteria and archaea. Nature biotechnology 35(8), 725–731 (2017)

[26] Olm, M.R., Crits-Christoph, A., Bouma-Gregson, K., Firek, B.A., Morowitz, M.J., Banfield, J.F.: instrain profiles population microdiversity from metagenomic data and sensitively detects shared microbial strains. Nature Biotechnology 39(6), 727–736 (2021)

[27] Fedarko, M.W., Martino, C., Morton, J.T., González, A., Rahman, G., Marotz, C.A., Minich, J.J., Allen, E.E., Knight, R.: Visualizing’omic feature rankings and log-ratios using qurro. NAR genomics and bioinformatics 2(2), 023 (2020)

[28] Morton, J.T., Marotz, C., Washburne, A., Silverman, J., Zaramela, L.S., Edlund, A., Zengler, K., Knight, R.: Establishing microbial composition measurement standards with reference frames. Nature communications 10(1), 2719 (2019)

[29] Tanno, H., Maeno, S., Salminen, S., Gueimonde, M., Endo, A.: 16s rrna gene sequence diversity in faecalibacterium prausnitzii-complex taxa has marked impacts on quantitative analysis. FEMS microbiology ecology 98(1), 004 (2022)

[30] Furusawa, Y., Obata, Y., Fukuda, S., Endo, T.A., Nakato, G., Takahashi, D., Nakanishi, Y., Uetake, C., Kato, K., Kato, T., et al.: Commensal microbe-derived butyrate induces the differentiation of colonic regulatory t cells. Nature 504(7480), 446–450 (2013)

[31] Arpaia, N., Campbell, C., Fan, X., Dikiy, S., Van Der Veeken, J., Deroos, P., Liu, H., Cross, J.R., Pfeffer, K., Coffer, P.J., et al.: Metabolites produced by commensal bacteria promote peripheral regulatory t-cell generation. Nature 504(7480), 451–455 (2013)

[32] Park, J., Kim, M., Kang, S.G., Jannasch, A.H., Cooper, B., Patterson, J., Kim, C.H.: Short-chain fatty acids induce both effector and regulatory t cells by suppression of histone deacetylases and regulation of the mtor–s6k pathway. Mucosal immunology 8(1), 80–93 (2015)

[33] Luu, M., Riester, Z., Baldrich, A., Reichardt, N., Yuille, S., Busetti, A., Klein, M., Wempe, A., Leister, H., Raifer, H., et al.: Microbial short-chain fatty acids modulate cd8+ t cell responses and improve adoptive immunotherapy for cancer. Nature communications 12(1), 4077 (2021)

[34] He, Y., Fu, L., Li, Y., Wang, W., Gong, M., Zhang, J., Dong, X., Huang, J., Wang, Q., Mackay, C.R., et al.: Gut microbial metabolites facilitate anticancer therapy efficacy by modulating cytotoxic cd8+ t cell immunity. Cell metabolism 33(5), 988–1000 (2021)

[35] Gloor, G.B., Macklaim, J.M., Pawlowsky-Glahn, V., Egozcue, J.J.: Microbiome datasets are compositional: and this is not optional. Frontiers in microbiology 8, 2224 (2017)

[36] Zhao, H., Li, D., Liu, J., Zhou, X., Han, J., Wang, L., Fan, Z., Feng, L., Zuo, J., Wang, Y.: Bifidobacterium breve predicts the efficacy of anti-pd-1 immunotherapy combined with chemotherapy in chinese nsclc patients. Cancer Medicine 12(5), 6325–6336 (2023)

[37] Lee, S.-H., Cho, S.-Y., Yoon, Y., Park, C., Sohn, J., Jeong, J.-J., Jeon, B.-N., Jang, M., An, C., Lee, S., et al.: Bifidobacterium bifidum strains synergize with immune checkpoint inhibitors to reduce tumour burden in mice. Nature Microbiology 6(3), 277–288 (2021)

[38] Yoon, Y., Kim, G., Jeon, B.-N., Fang, S., Park, H.: Bifidobacterium strain-specific enhances the efficacy of cancer therapeutics in tumor-bearing mice. Cancers 13(5), 957 (2021)

[39] Daill’ere, R., Vétizou, M., Waldschmitt, N., Yamazaki, T., Isnard, C., Poirier-Colame, V., Duong, C.P., Flament, C., Lepage, P., Roberti, M.P., et al.: Enterococcus hirae and barnesiella intestinihominis facilitate cyclophosphamide-induced therapeutic immunomodulatory effects. Immunity 45(4), 931–943 (2016)

[40] Dizman, N., Hsu, J., Bergerot, P.G., Gillece, J.D., Folkerts, M., Reining, L., Trent, J., Highlander, S.K., Pal, S.K.: Randomized trial assessing impact of probiotic supplementation on gut microbiome and clinical outcome from targeted therapy in metastatic renal cell carcinoma. Cancer Medicine 10(1), 79–86 (2021)

[41] Mao, J., Wang, D., Long, J., Yang, X., Lin, J., Song, Y., Xie, F., Xun, Z., Wang, Y., Wang, Y., et al.: Gut microbiome is associated with the clinical response to anti-pd-1 based immunotherapy in hepatobiliary cancers. Journal for Immunotherapy of Cancer 9(12) (2021)

[42] Peters, B.A., Wilson, M., Moran, U., Pavlick, A., Izsak, A., Wechter, T., Weber, J.S., Osman, I., Ahn, J.: Relating the gut metagenome and metatranscriptome to immunotherapy responses in melanoma patients. Genome medicine 11, 1–14 (2019)

[43] Bachem, A., Makhlouf, C., Binger, K.J., Souza, D.P., Tull, D., Hochheiser, K., Whitney, P.G., Fernandez-Ruiz, D., Dähling, S., Kastenmüller, W., et al.: Microbiota-derived short-chain fatty acids promote the memory potential of antigen-activated cd8+ t cells. Immunity 51(2), 285–297 (2019)

[44] Danne, C., Sokol, H.: Butyrate, a new microbiota-dependent player in cd8+ t cells immunity and cancer therapy? Cell Reports Medicine 2(7) (2021)

[45] Rios-Covian, D., Gueimonde, M., Duncan, S.H., Flint, H.J., Los Reyes-Gavilan, C.G.: Enhanced butyrate formation by cross-feeding between faecalibacterium prausnitzii and bifidobacterium adolescentis. FEMS microbiology letters 362(21), 176 (2015)

[46] Kim, H., Jeong, Y., Kang, S., You, H.J., Ji, G.E.: Co-culture with bifidobacterium catenulatum improves the growth, gut colonization, and butyrate production of faecalibacterium prausnitzii: in vitro and in vivo studies. Microorganisms 8(5), 788 (2020)

[47] Lapébie, P., Lombard, V., Drula, E., Terrapon, N., Henrissat, B.: Bacteroidetes use thousands of enzyme combinations to break down glycans. Nature communications 10(1), 2043 (2019)

[48] Feng, J., Qian, Y., Zhou, Z., Ertmer, S., Vivas, E.I., Lan, F., Hamilton, J.J., Rey, F.E., Anantharaman, K., Venturelli, O.S.: Polysaccharide utilization loci in bacteroides determine population fitness and community-level interactions. Cell host & microbe 30(2), 200–215 (2022)

[49] Degnan, P.H., Taga, M.E., Goodman, A.L.: Vitamin b12 as a modulator of gut microbial ecology. Cell metabolism 20(5), 769–778 (2014)

[50] Juodeikis, R., Jones, E., Deery, E., Beal, D.M., Stentz, R., Kräutler, B., Carding, S.R., Warren, M.J.: Nutrient smuggling: Commensal gut bacteria-derived extracellular vesicles scavenge vitamin b12 and related cobamides for microbe and host acquisition. Journal of Extracellular Biology 1(10), 61 (2022)

[51] Plichta, D.R., Juncker, A.S., Bertalan, M., Rettedal, E., Gautier, L., Varela, E., Manichanh, C., Fouqueray, C., Levenez, F., Nielsen, T., et al.: Transcriptional interactions suggest niche segregation among microorganisms in the human gut. Nature Microbiology 1(11), 1–6 (2016)

[52] Koh, A., De Vadder, F., Kovatcheva-Datchary, P., Bäckhed, F.: From dietary fiber to host physiology: short-chain fatty acids as key bacterial metabolites. Cell 165(6), 1332–1345 (2016)

[53] Jugder, B.-E., Kamareddine, L., Watnick, P.I.: Microbiota-derived acetate activates intestinal innate immunity via the tip60 histone acetyltransferase complex. Immunity 54(8), 1683–1697 (2021)

[54] Yu, T., Yang, W., Yao, S., Yu, Y., Wakamiya, M., Golovko, G., Cong, Y.: Sting promotes intestinal iga production by regulating acetate-producing bacteria to maintain host-microbiota mutualism. Inflammatory Bowel Diseases 29(6), 946– 959 (2023)

[55] Barnes, R.H., Fiala, G.: Effects of the prevention of coprophagy in the rat: Ii. vitamin b12 requirement. The Journal of Nutrition 65(1), 103–114 (1958)

[56] Morgan, T., Gregory, M.E., Kon, S., Porter, J.: Coprophagy and vitamin b12 in the rat. British Journal of Nutrition 18(1), 595–602 (1964)

[57] Si, W., Liang, H., Bugno, J., Xu, Q., Ding, X., Yang, K., Fu, Y., Weichselbaum, R.R., Zhao, X., Wang, L.: Lactobacillus rhamnosus gg induces cgas/sting-dependent type i interferon and improves response to immune checkpoint blockade. Gut 71(3), 521–533 (2022)

[58] Gao, G., Shen, S., Zhang, T., Zhang, J., Huang, S., Sun, Z., Zhang, H.: Lactica-seibacillus rhamnosus probio-m9 enhanced the antitumor response to anti-pd-1 therapy by modulating intestinal metabolites. Ebiomedicine 91 (2023)

[59] Dizman, N., Meza, L., Bergerot, P., Alcantara, M., Dorff, T., Lyou, Y., Frankel, P., Cui, Y., Mira, V., Llamas, M., et al.: Nivolumab plus ipilimumab with or without live bacterial supplementation in metastatic renal cell carcinoma: a randomized phase 1 trial. Nature medicine 28(4), 704–712 (2022)

[60] Khan, M.T., Dwibedi, C., Sundh, D., Pradhan, M., Kraft, J.D., Caesar, R., Tremaroli, V., Lorentzon, M., Bäckhed, F.: Synergy and oxygen adaptation for development of next-generation probiotics. Nature, 1–5 (2023)

[61] Bolger, A.M., Lohse, M., Usadel, B.: Trimmomatic: a flexible trimmer for illumina sequence data. Bioinformatics 30(15), 2114–2120 (2014)

[62] Bushnell, B.: Bbmap: a fast, accurate, splice-aware aligner. Technical report, Lawrence Berkeley National Lab.(LBNL), Berkeley, CA (United States) (2014)

[63] Li, D., Liu, C.-M., Luo, R., Sadakane, K., Lam, T.-W.: Megahit: an ultra-fast single-node solution for large and complex metagenomics assembly via succinct de bruijn graph. Bioinformatics 31(10), 1674–1676 (2015)

[64] Wu, Y.-W., Simmons, B.A., Singer, S.W.: Maxbin 2.0: an automated binning algorithm to recover genomes from multiple metagenomic datasets. Bioinformatics 32(4), 605–607 (2016)

[65] Kang, D.D., Li, F., Kirton, E., Thomas, A., Egan, R., An, H., Wang, Z.: Metabat 2: an adaptive binning algorithm for robust and efficient genome reconstruction from metagenome assemblies. PeerJ 7, 7359 (2019)

[66] Sieber, C.M., Probst, A.J., Sharrar, A., Thomas, B.C., Hess, M., Tringe, S.G., Banfield, J.F.: Recovery of genomes from metagenomes via a dereplication, aggregation and scoring strategy. Nature microbiology 3(7), 836–843 (2018)

[67] Olm, M.R., Brown, C.T., Brooks, B., Banfield, J.F.: drep: a tool for fast and accurate genomic comparisons that enables improved genome recovery from metagenomes through de-replication. The ISME journal 11(12), 2864–2868 (2017)

[68] Parks, D.H., Imelfort, M., Skennerton, C.T., Hugenholtz, P., Tyson, G.W.: Checkm: assessing the quality of microbial genomes recovered from isolates, single cells, and metagenomes. Genome research 25(7), 1043–1055 (2015)

[69] Chaumeil, P.-A., Mussig, A.J., Hugenholtz, P., Parks, D.H.: Gtdb-tk v2: memory friendly classification with the genome taxonomy database. Bioinformatics 38(23), 5315–5316 (2022)

[70] Parks, D.H., Chuvochina, M., Rinke, C., Mussig, A.J., Chaumeil, P.-A., Hugenholtz, P.: Gtdb: an ongoing census of bacterial and archaeal diversity through a phylogenetically consistent, rank normalized and complete genome-based taxonomy. Nucleic acids research 50(D1), 785–794 (2022)

[71] Price, M.N., Dehal, P.S., Arkin, A.P.: Fasttree 2–approximately maximum-likelihood trees for large alignments. PloS one 5(3), 9490 (2010)

[72] Edgar, R.C.: Muscle: a multiple sequence alignment method with reduced time and space complexity. BMC bioinformatics 5(1), 1–19 (2004)

[73] Cantrell, K., Fedarko, M.W., Rahman, G., McDonald, D., Yang, Y., Zaw, T., Gonzalez, A., Janssen, S., Estaki, M., Haiminen, N., et al.: Empress enables tree-guided, interactive, and exploratory analyses of multi-omic data sets. MSystems 6(2), 01216–20 (2021)

[74] Bolyen, E., Rideout, J.R., Dillon, M.R., Bokulich, N.A., Abnet, C.C., Al-Ghalith, G.A., Alexander, H., Alm, E.J., Arumugam, M., Asnicar, F., et al.: Reproducible, interactive, scalable and extensible microbiome data science using qiime 2. Nature biotechnology 37(8), 852–857 (2019)

[75] Martino, C., Morton, J.T., Marotz, C.A., Thompson, L.R., Tripathi, A., Knight, R., Zengler, K.: A novel sparse compositional technique reveals microbial perturbations. MSystems 4(1), 10–1128 (2019)

[76] Hyatt, D., Chen, G.-L., LoCascio, P.F., Land, M.L., Larimer, F.W., Hauser, L.J.: Prodigal: prokaryotic gene recognition and translation initiation site identification. BMC bioinformatics 11, 1–11 (2010)

[77] Emms, D.M., Kelly, S.: Orthofinder: phylogenetic orthology inference for comparative genomics. Genome biology 20, 1–14 (2019)

[78] Xu, S., Li, L., Luo, X., Chen, M., Tang, W., Zhan, L., Dai, Z., Lam, T.T., Guan, Y., Yu, G.: Ggtree: a serialized data object for visualization of a phylogenetic tree and annotation data. IMeta 1(4), 56 (2022)

[79] Orakov, A., Fullam, A., Coelho, L.P., Khedkar, S., Szklarczyk, D., Mende, D.R., Schmidt, T.S., Bork, P.: Gunc: detection of chimerism and contamination in prokaryotic genomes. Genome biology 22, 1–19 (2021)

[80] Drula, E., Garron, M.-L., Dogan, S., Lombard, V., Henrissat, B., Terrapon, N.: The carbohydrate-active enzyme database: functions and literature. Nucleic acids research 50(D1), 571–577 (2022)

[81] Kanehisa, M., Furumichi, M., Tanabe, M., Sato, Y., Morishima, K.: Kegg: new perspectives on genomes, pathways, diseases and drugs. Nucleic acids research 45(D1), 353–361 (2017)

[82] Buchfink, B., Reuter, K., Drost, H.-G.: Sensitive protein alignments at tree-of-life scale using diamond. Nature methods 18(4), 366–368 (2021)

[83] Zimmermann, J., Kaleta, C., Waschina, S.: gapseq: Informed prediction of bacterial metabolic pathways and reconstruction of accurate metabolic models. Genome biology 22(1), 1–35 (2021)

[84] Caspi, R., Billington, R., Fulcher, C.A., Keseler, I.M., Kothari, A., Krummenacker, M., Latendresse, M., Midford, P.E., Ong, Q., Ong, W.K., et al.: The metacyc database of metabolic pathways and enzymes. Nucleic acids research 46(D1), 633–639 (2018)

[85] Oksanen, J.: Vegan: community ecology package. http://vegan.r-forge.r-project.org/ (2010)

[86] Väremo, L., Nielsen, J., Nookaew, I.: Enriching the gene set analysis of genome-wide data by incorporating directionality of gene expression and combining statistical hypotheses and methods. Nucleic acids research 41(8), 4378–4391 (2013)

[87] Zhang, B., Lingga, C., Bowman, C., Hackmann, T.J.: A new pathway for forming acetate and synthesizing atp during fermentation in bacteria. Applied and environmental microbiology 87(14), 02959–20 (2021)

[88] Esposito, A., Tamburini, S., Triboli, L., Ambrosino, L., Chiusano, M.L., Jousson, O.: Insights into the genome structure of four acetogenic bacteria with specific reference to the wood–ljungdahl pathway. Microbiologyopen 8(12), 938 (2019)

[89] Vital, M., Howe, A.C., Tiedje, J.M.: Revealing the bacterial butyrate synthesis pathways by analyzing (meta) genomic data. MBio 5(2), 10–1128 (2014)

[90] Frolova, M.S., Suvorova, I.A., Iablokov, S.N., Petrov, S.N., Rodionov, D.A.: Genomic reconstruction of short-chain fatty acid production by the human gut microbiota. Frontiers in Molecular Biosciences 9, 949563 (2022)

[91] Gelius-Dietrich, G., Desouki, A.A., Fritzemeier, C.J., Lercher, M.J.: Sybil– efficient constraint-based modelling in r. BMC systems biology 7, 1–8 (2013)

