## Supplementary figures and images for "Systemic Metabolic Depletion of Intestine Microbiome Undermines Melanoma Immunotherapy Effectiveness"

### Figure_S1.pdf

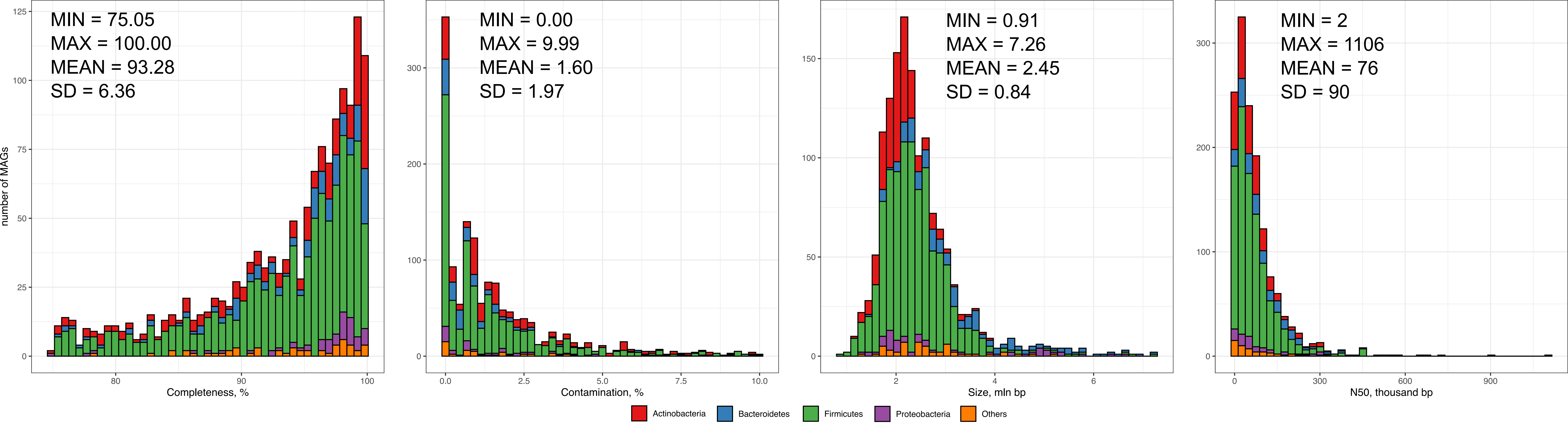

### Figure_S2.pdf

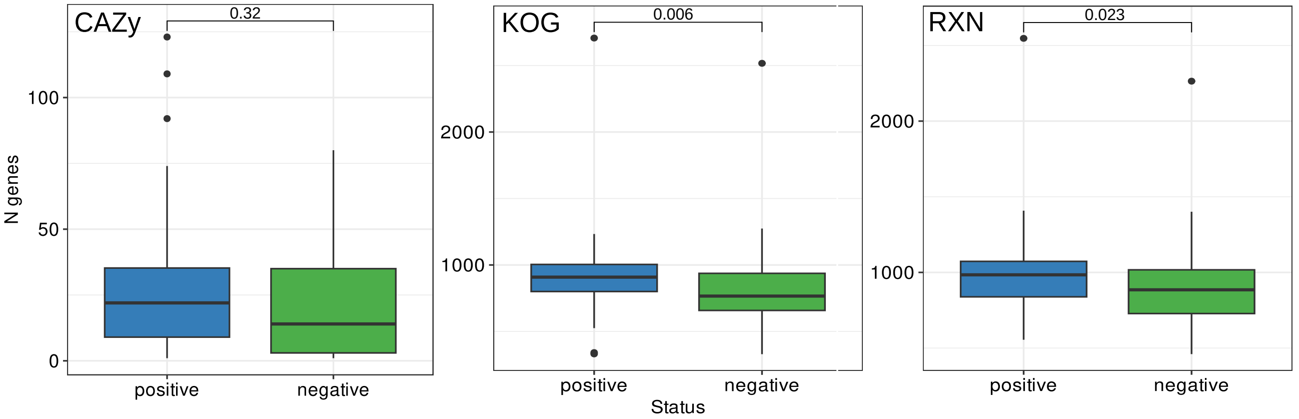

### Figure_S3.pdf

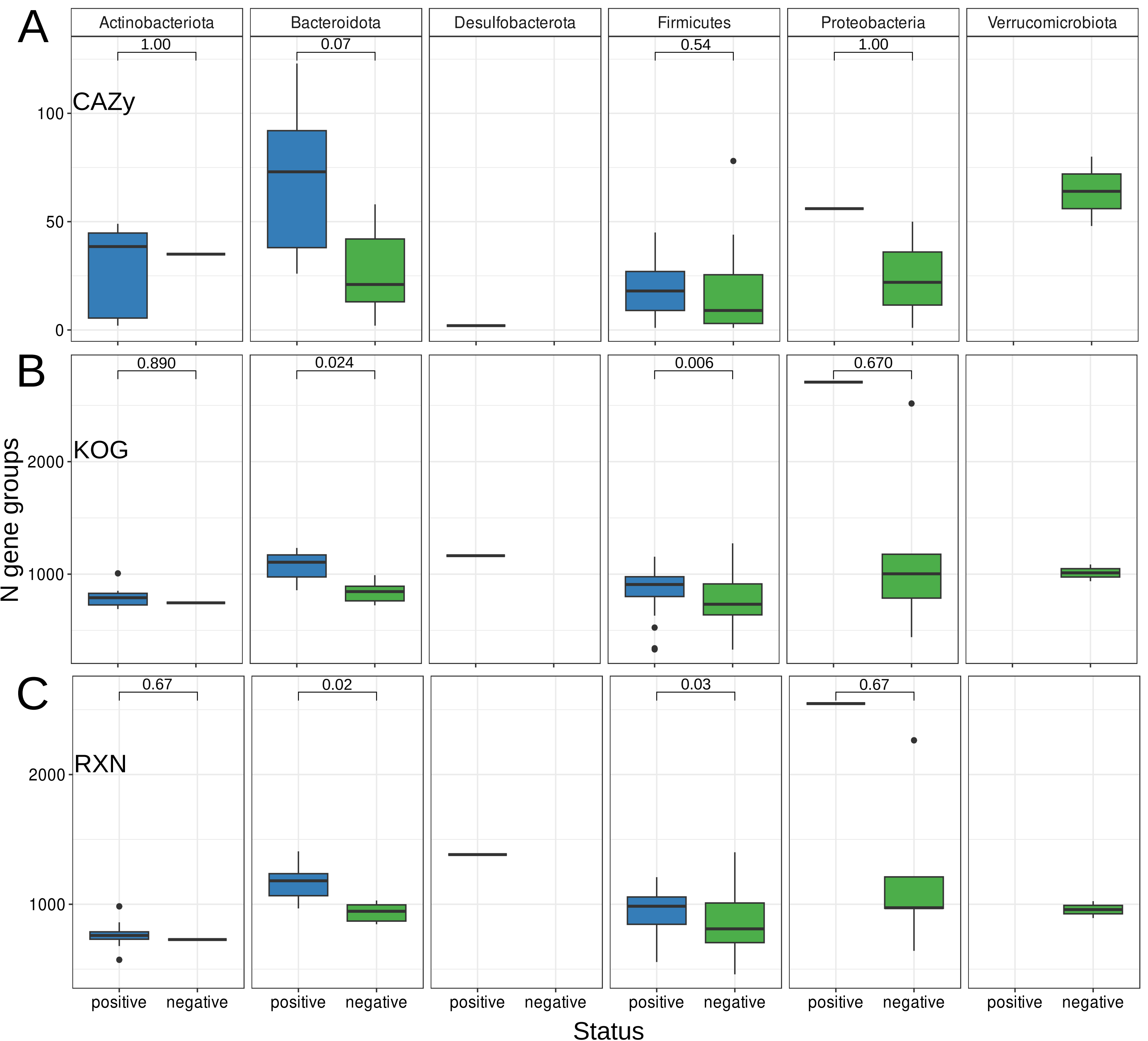

### Figure_S4.pdf

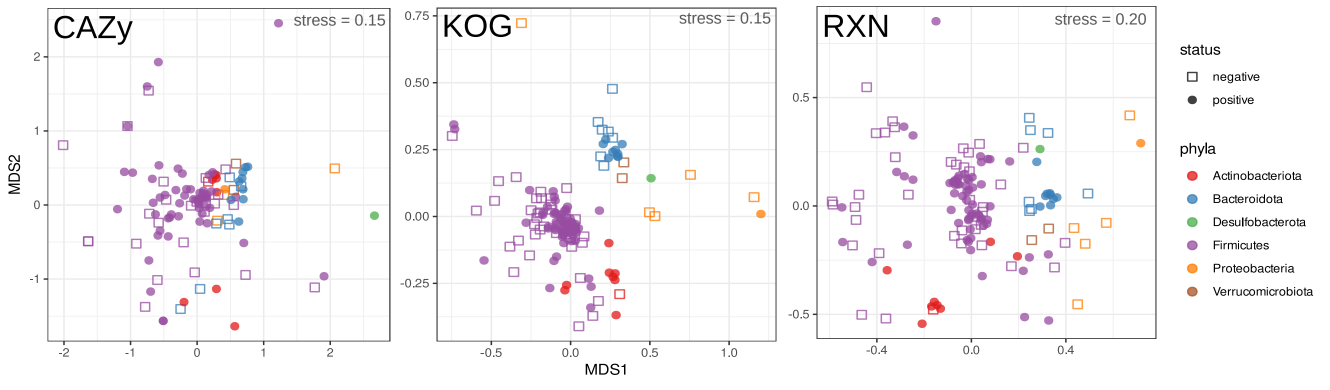

### Figure_S5.png

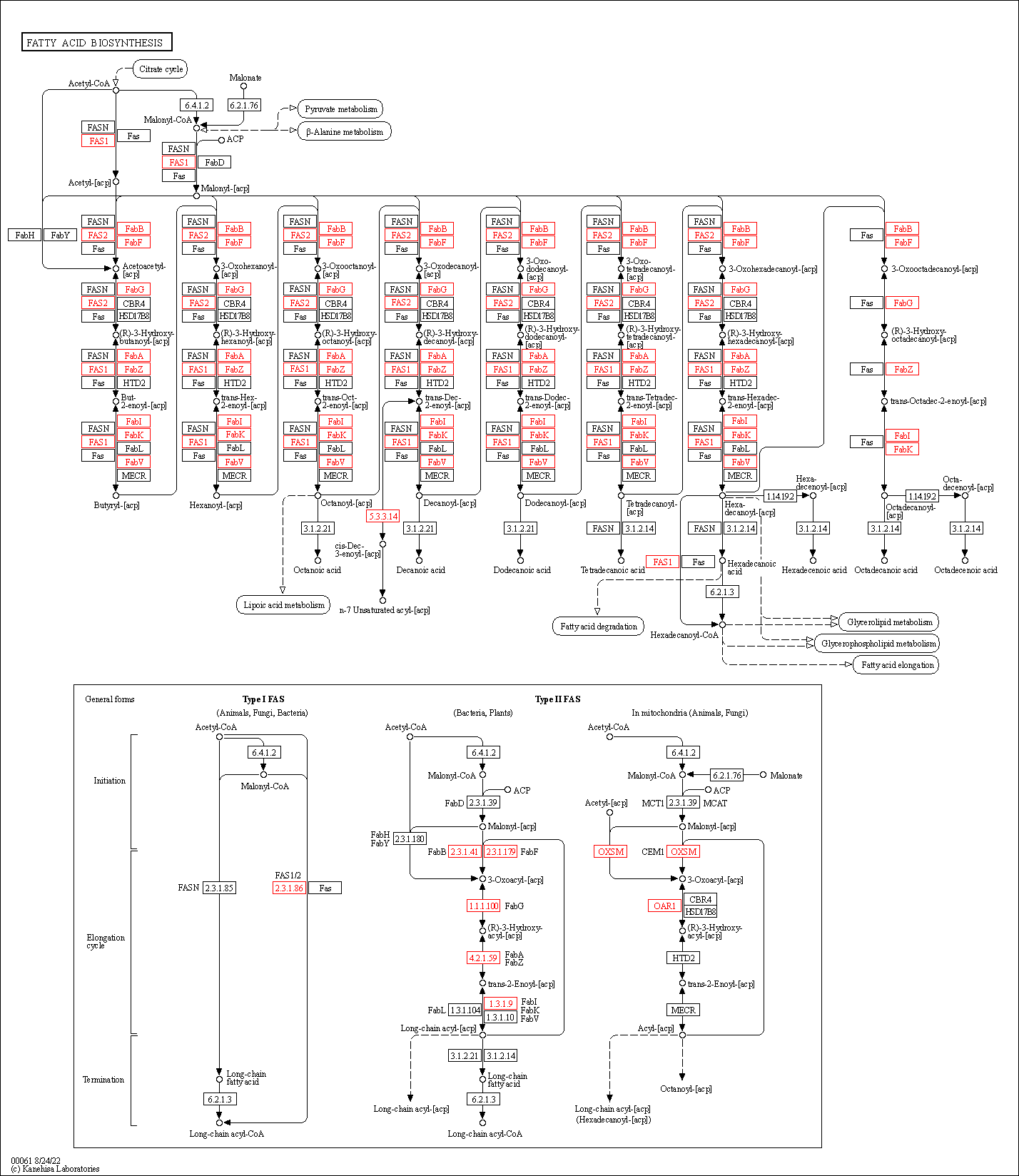

### Figure_S6.pdf

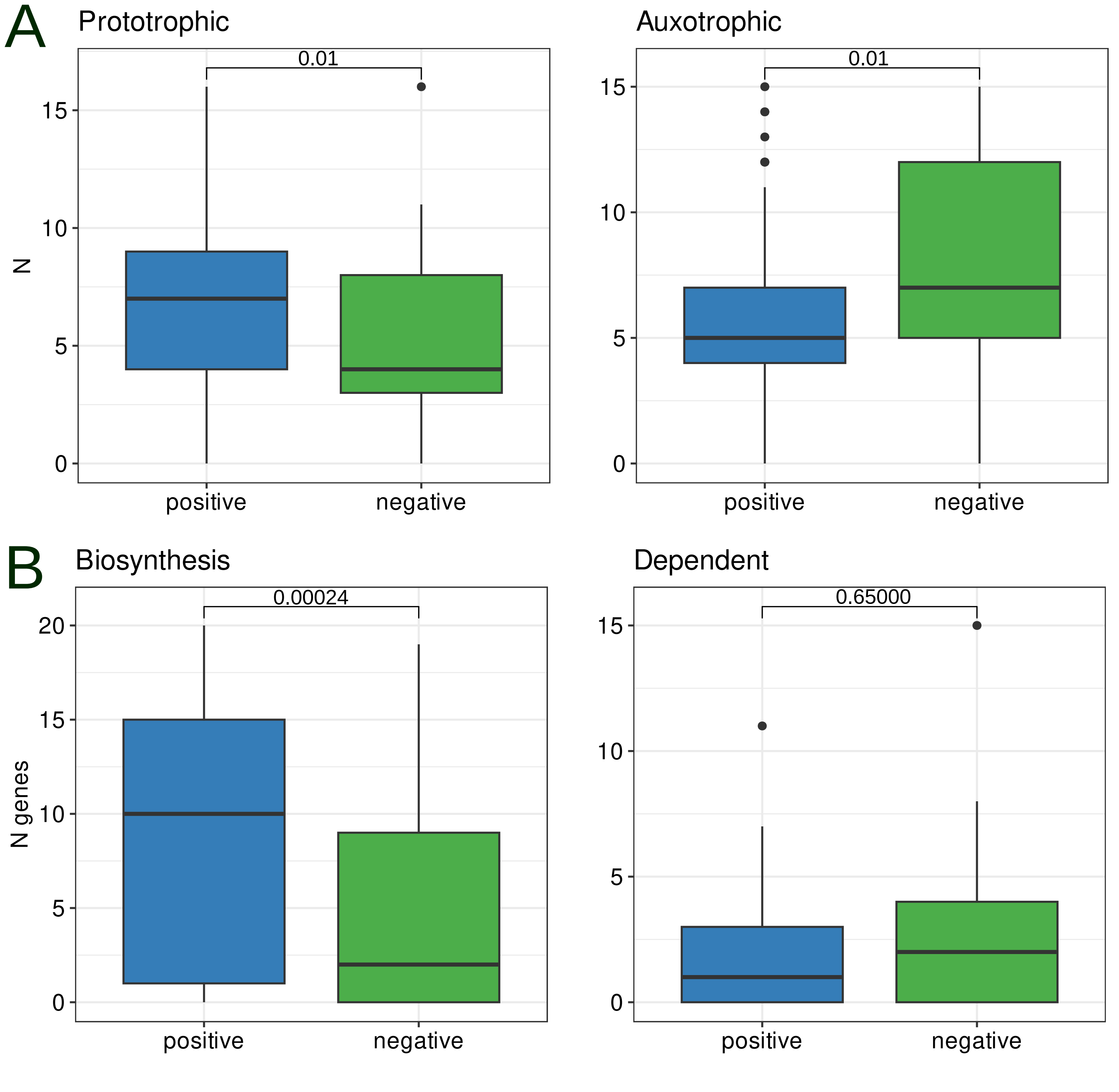

### Figure_S7.png

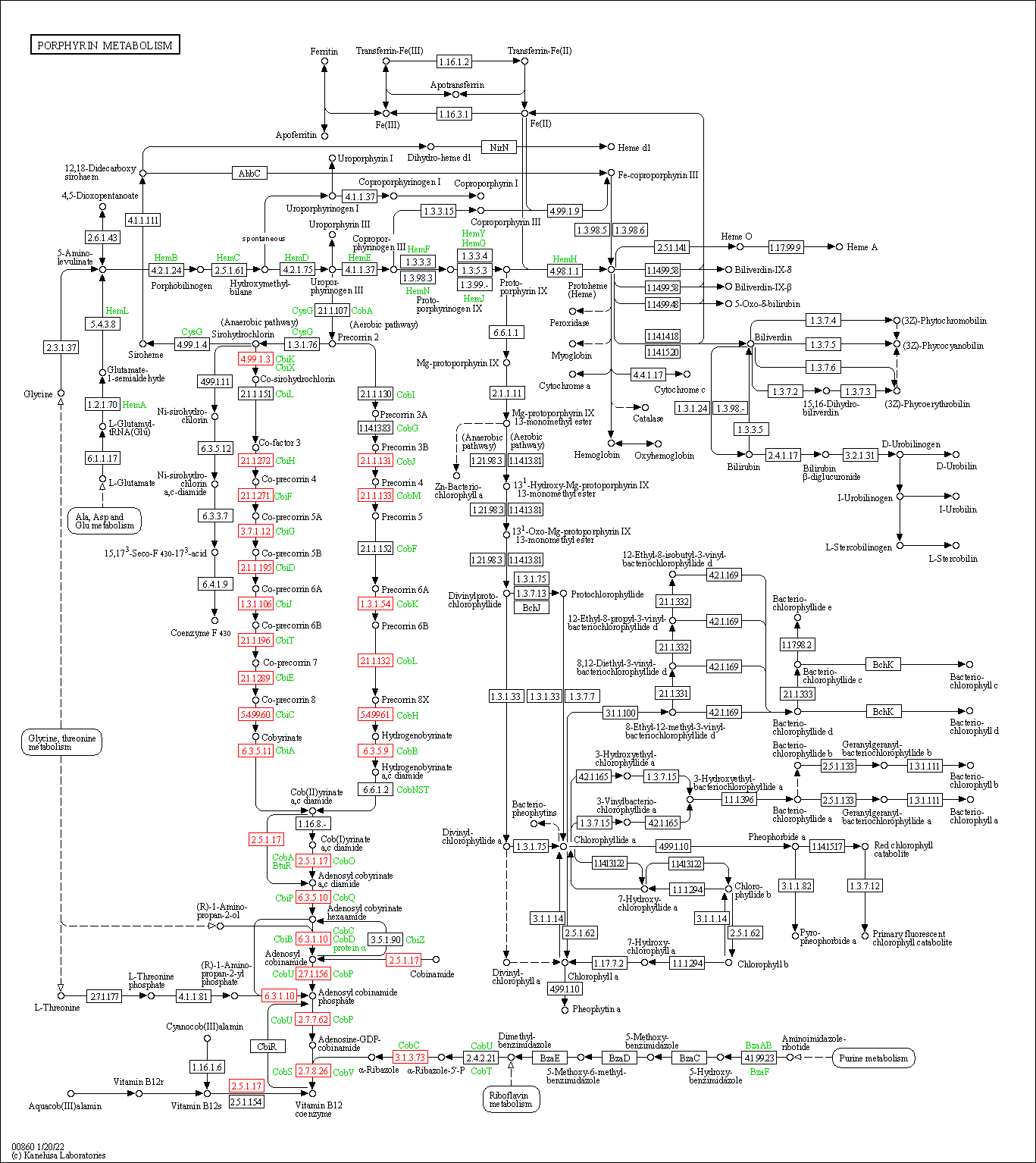

### Figure_S8.pdf

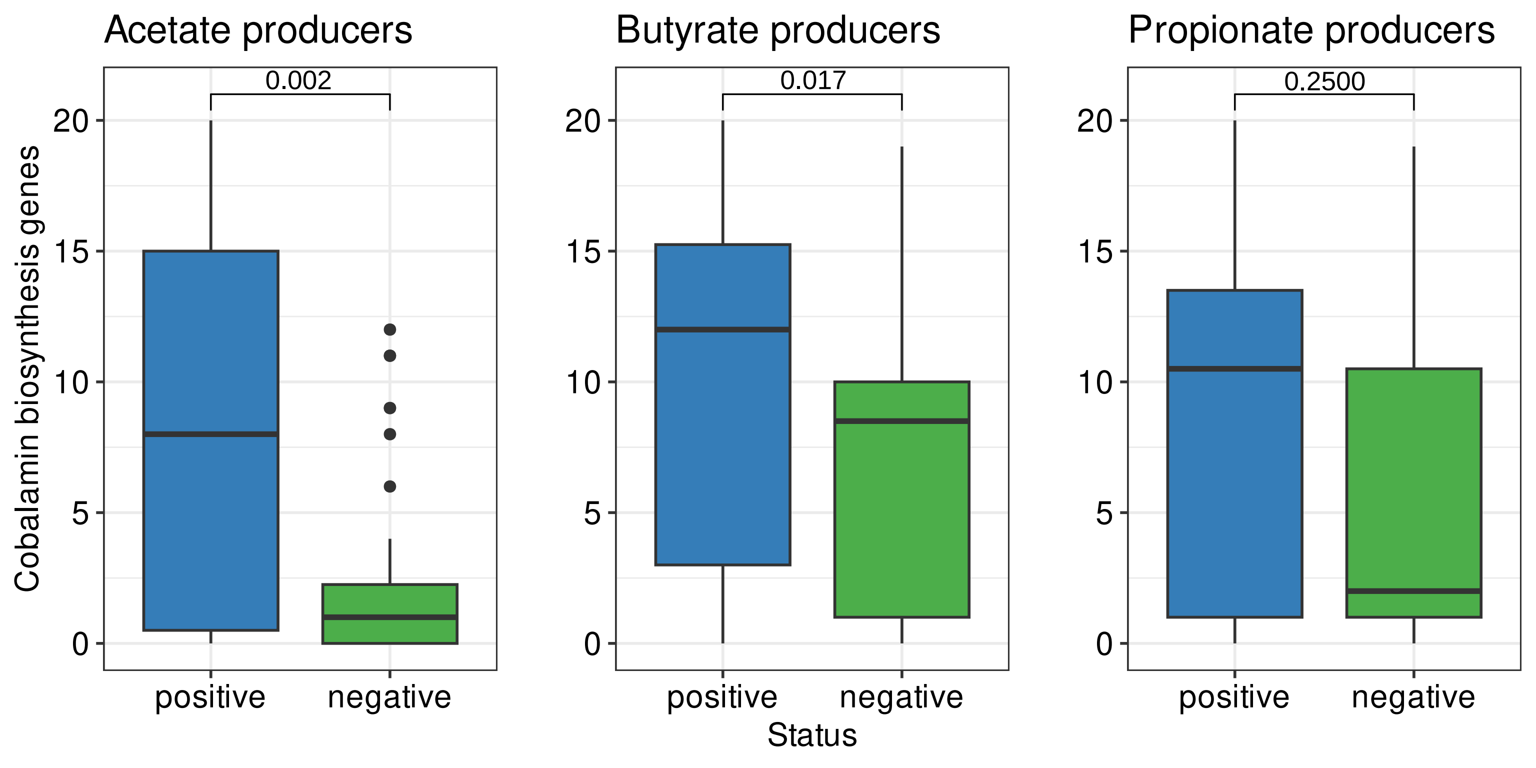
